# A Review and Evaluation of Species Richness Estimation

**DOI:** 10.1101/2024.10.09.615408

**Authors:** Johanna Elena Schmitz, Sven Rahmann

## Abstract

**Motivation:** The statistical problem of estimating the total number of distinct species in a population (or distinct elements in a multiset), given only a small sample, occurs in various areas, ranging from the unseen species problem in ecology to estimating the diversity of immune repertoires. Accurately estimating the true richness from very small samples is challenging, in particular for highly diverse populations with many rare species. Depending on the application, different estimation strategies have been proposed that incorporate explicit or implicit assumptions about either the species distribution or about the sampling process. These methods are scattered across the literature, and an extensive overview of their assumptions, methodology and performance is currently lacking.

**Results:** We comprehensively review and evaluate a variety of existing methods on real and simulated data with different compositions of rare and abundant elements. Our evaluation shows that, depending on species composition, different methods provide the most accurate richness estimates. Simpler methods, like the Chao 1 and Chiu estimators, yield accurate predictions for many of the tested species compositions, but tend to underestimate the true richness for heterogeneous populations and small (containing 1% to 5% of the population) samples. When the population size is known, upsampling estimators such as PreSeq and RichnEst often yield more accurate results.

**Availability and implementation:** Source code for data simulation and richness estimation is available at https://gitlab.com/rahmannlab/speciesrichness.

## 1 Introduction

Estimating the diversity of a population from a small sample has a wide range of applications in diverse fields, such as ecology, immunology, biological sequence analysis and linguistics. One of the oldest applications is the unseen species problem in ecology (Fisher et al., 1943), e.g., predicting the number of butterfly species on an island after capturing a small collection of butterflies. The same statistical problem arises in linguistics when trying to estimate how many words a writer might have known but never used in any of his published works, which is a measure in quantitative linguistics to compare the vocabulary richness of writers (Good, 1953). Recent applications include the analysis of microbial complexity in environmental niches (Datta et al., 2020), the comparison of bacterial diversity in human guts under different disease conditions (Hills et al., 2019), or the quantification of a suitable sequencing depth to study rare cancer types based on the diversity of genetic variants and mutations (Masoero et al., 2022).

While there are many measures of diversity, such as the proportion of rare and abundant species, or the entropy of the species distribution, we limit this review to the estimation of *species richness*, which measures the total number of distinct species in a population, assuming that each individual belongs to a single species. We hence exclude methods that assume that an individual may belong to several classes at once.

In most applications, it is infeasible to observe the complete population; so, the *observed* species richness of a sample usually underestimates the *true* richness, especially for populations with many rare species. However, an accurate estimate is crucial to analyse the properties of a population. For instance, the T- or B-cell receptor richness of immune repertoires indicates the effectiveness of the immune system, and an accurate estimate is thus vital to compare immune systems between healthy and diseased individuals. Since the frequency distribution of T-cell receptor repertoires is highly skewed, rare T-cell receptors are often missed in the sampling process which necessitates robust estimation of the true richness (Laydon et al., 2015).

Accurate estimation of species richness is a challenging statistical problem, in particular without making additional assumptions about the sampling process or the species distribution (see Figure 1). Various estimators have been proposed over the years to achieve accurate richness estimates for populations with different species compositions. Early estimators, like the Chao 1 (Chao, 1984) or Jackknife estimator (Burnham and Overton, 1978), assume that most information about the number of missing species is present in the number of species captured only once or twice. Other estimators assume that the species counts follow a parametric probability distribution, e.g., a Gamma-Poisson mixture distribution (Sandland and Cormack, 1984). Several recent methods make no such assumptions and are based on linear programming (Schröder and Rahmann, 2015; Valiant and Valiant, 2017) or curve fitting Willis and Bunge (2015); Laydon et al. (2015).

**Figure 1.**
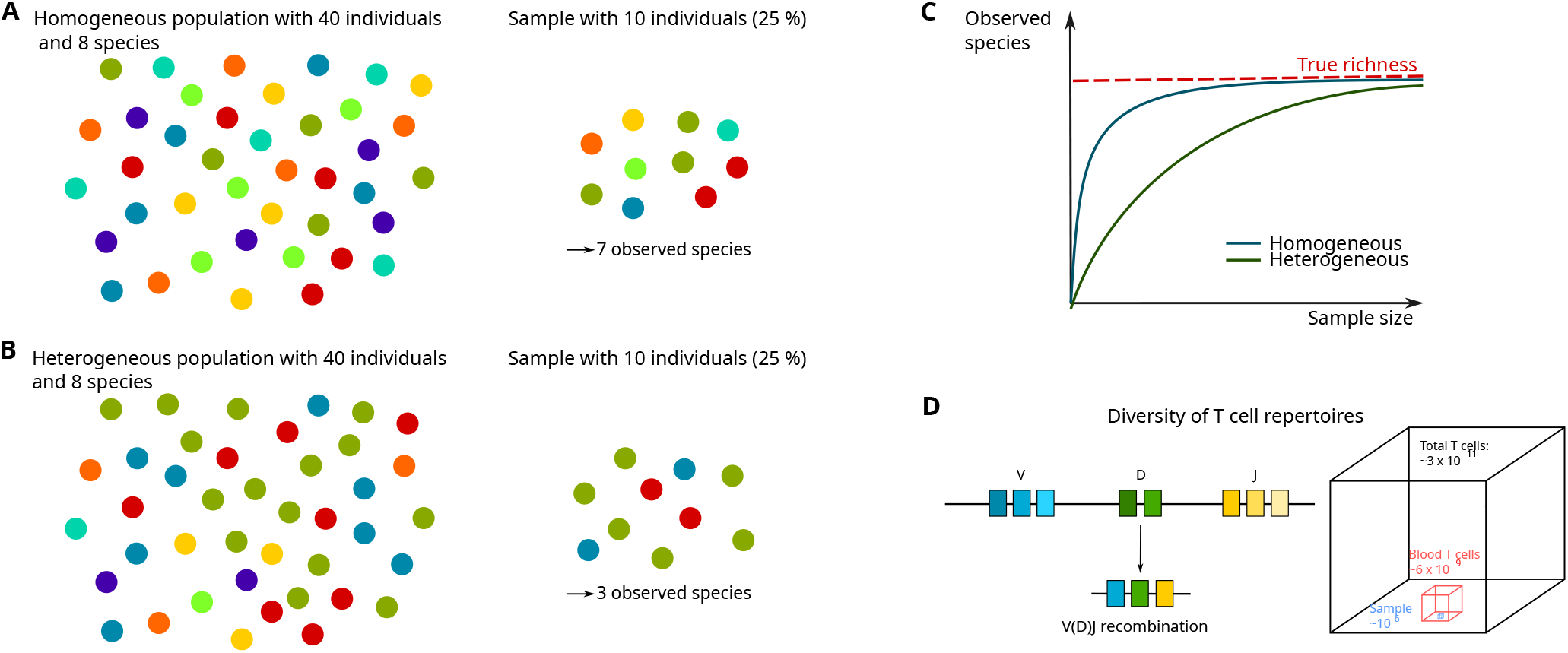
Comparison of species (color) richness in the population and in a sample. **A** For a homogeneous population, a small sample is sufficient to observe a sample richness that is close to the true richness. **B** The sample richness of a population with many rare species underestimates the true richness. **C** Rarefaction curve. Increasing the sample size leads to an increase in the observed richness, converging to the true richness. For a heterogeneous population with many rare species, the convergence is slow. **D** T-cell diversity. Somatic recombination of V, D, J gene segments during T-cell maturation gives rise to numerous distinct T-cell receptors that form an immune system that is able to recognize almost all potential pathogens. However, analyzing T-cell receptor diversity based on a small blood sample is challenging, because the sample only contains a minute portion of all T-cells from a person.

Since a systematic comparison of long-established and contemporary species richness estimators has not yet been conducted, we evaluate the performance of species richness estimators on a variety of simulated and real data with different underlying frequency distributions.

## 2 Methods

### 2.1 Definitions and Notation

From a full population of *N* individuals (*N* may be finite or infinite, known or unknown), a finite random sample of *n* individuals is observed. We assume that the sample is small, i.e., *n* ≪ *N*. Each individual (sometimes called element) in the population belongs to a species (sometimes called class or group). The number of species in the full population is referred to as its *species richness S*, which we assume to be finite (even for infinite *N*).

The *observed sample richness* is denoted by *S*_obs_.

For each observed species, we count its abundance in the sample and obtain the *abundance* vector 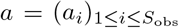.

The number of species observed exactly *k* times in the sample is given by *f*_*k*_ (i.e., *f*_*k*_ = |{*i* | *a*_*i*_ = *k*, 1 ≤ *i* ≤ *S*_obs_}|), such that the number *n* of individuals in the sample satisfies

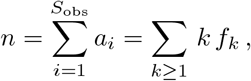

and the observed richness is given by

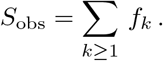

The population’s species richness can be expressed by

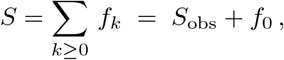

i.e., the observed richness plus the number of unobserved species that are missing in the sample. Therefore, estimating *S* is equivalent to estimating the unobserved *f*_0_ from the observed *f* = (*f*_1_, *f*_2_, …). As the observation is finite, *f* is of finite length.

### 2.2 Classification of Richness Estimators

Species richness estimators can be divided into two main groups: (1) If the total population size *N* is known or assumed to be known, finite, and given as an input, we have an *upsampling* task (by a factor of *N/n*), i.e., we need to solve the inverse problem of the random (down)sampling process. (2) If *N* is unknown or assumed infinite, it is (often implicitly) assumed that the total species richness *S* is finite, i.e., the rarefaction curve reaches a finite asymptotic upper limit (Fig. 1C), which means that there cannot be arbitrarily many rare species. The first group is referred to as *upsampling estimators* and the second group as *population estimators*. If (an approximation of) *N* is available, an upsampling estimator should be preferred, as using more information typically yields more accurate results.

#### Scenarios for upsampling and population estimators

A use case for an upsampling estimator is deciding about the sequencing depth of a DNA library of unknown quality. Based on PCR duplicate statistics after a low-depth (say, 3x coverage) pre-experiment, it can be decided whether 30x vs. 15x coverage would yield significantly more new fragments, or whether one would see mostly PCR duplicates. A use case for both classes is the estimation of the richness of T-cell receptor repertoires from small blood samples (e.g., in healthy vs. sick individuals): The exact total number of T-cells is unknown, but we have reasonable estimates of the total number of T-cells in the body and in the blood (Fig. 1D). A use case for population estimators is to estimate the microbiome species diversity of the intestine or to estimate the bacterial species diversity in soil.

#### Classification of estimators by assumptions made

Estimators can be further divided according to whether they make assumptions about the species composition, i.e., about the behavior of (*f*_*k*_)_*k*≥0_. If they do, the species richness estimation problem often simplifies to estimating one or a few parameters of a parametric distribution, which leads to computationally efficient estimators that show good accuracy if the assumptions are satisfied, but that may be inaccurate if not. We call these estimators *parametric estimators*, and estimators that make no explicit distributional assumptions *non-parametric estimators*.

An overview of estimators that we discuss in more detail in the following sections appears in Table 1. Each particular method may make explicit or implicit additional assumptions, which we shall describe below as needed. We first mention several common principles behind these methods.

**Table 1:** Overview of species richness estimators. Column “up” indicates whether an estimator is an upsampling estimator 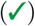 or not (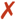; then it is a population estimator). Population and upsampling estimators are separated by a horizontal line. Column “par” indicates whether an estimator is parametric 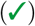, i.e., whether it makes distributional assumptions about the species composition, or not 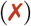. Column “Implementation” links to the estimator’s implementation that we used for the computational experiments (“ours” means our own implementation at https://gitlab.com/rahmannlab/speciesrichness).

| Name | up | par | Reference | Implementation |
| --- | --- | --- | --- | --- |
| Good-Turing | ✗ | ✗ | Good (1953) | ours |
| Jackknife | ✗ | ✗ | Burnham and Overton (1978) | ours |
| ACE | ✗ | ✗ | Chao and Lee (1992) | ours |
| Poisson | ✗ | ✓ | Sandland and Cormack (1984) | Breakaway R package |
| Chao 1 | ✗ | ✗✓* | Chao (1984) | ours |
| Gamma-Poisson mixture | ✗ | ✓ | Fisher et al. (1943) | ours |
| Chao and Bunge | ✗ | ✓ | Chao and Bunge (2002) | Breakaway R package |
| Lanumteang and Böhning | ✗ | ✓ | Lanumteang and Böhning (2011) | ours |
| Chiu | ✗ | ✓ | Chiu (2023) | ours |
| Objective Bayesian | ✗ | ✓ | Barger and Bunge (2010) | Breakaway R package |
| Recon | ✗ | ✗ | Kaplinsky and Arnaout (2016) | GitHub ArnaoutLab |
| Valiant | ✗ | ✗ | Valiant and Valiant (2017) | Valiant Code |
| Breakaway | ✗ | ✗ | Willis and Bunge (2015) | Breakaway R package |
| Smoothed Good-Toulmin | ✓ | ✓ | Orlitsky et al. (2016) | ours |
| PreSeq | ✓ | ✗ | Daley and Smith (2013) | PreSeq R package |
| Pitman sampling formula | ✓ | ✓ | Pitman (1995) | GitHub Stefanie Tauber |
| DivE | ✓ | ✗ | Laydon et al. (2014) | DivE R package |
| RichnEst (formerly Dupre) | ✓ | ✗ | Schröder and Rahmann (2015) | GitLab RahmannLab |

\* The Chao 1 estimator can be derived from a Poisson model, but was first introduced as a nonparametric estimator.

### 2.3 General Principles

Given the abundance vector *a* = (*a*_*i*_), *i* = 1, …, *S*_obs_ of the observed species, we assume that it is the realization of an underlying probabilistic model. Let *X* denote the random variable describing the abundance of a randomly picked species; let *p*_*k*_ be the probability of observing a species exactly *k* times; so ℙ[*X* = *k*] = *p*_*k*_ for *k* = 0, 1, 2, If we draw *S* times an independent copy of *X* (abundances, including zeros) and count how many times each abundance *k* was seen, we obtain *f* = (*f*_*k*_), including *f*_0_. Conversely, *f*_*k*_*/S* is an estimate for *p*_*k*_.

#### Zero-truncated distribution

The number of unobserved species *f*_0_ is unknown. Thus, if *P* = (*p*_*k*_)_*k*≥0_ with *p*_*k*_ being the true probability distribution for capturing a species exactly *k* times (whether it follows a parametric family or not), then 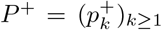 with 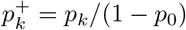 for *k* ≥ 1 is the *zero-truncated* distribution. It is obtained from *P* by setting 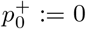 and 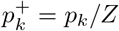, where the normalization constant *Z* ensures that 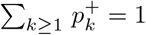mso *Z* = 1 − *p*_0_ = *S*_obs_*/S*. From this, we derive

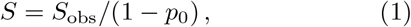

which by itself is not very helpful, as both *S* and *p*_0_ are unknown, but with additional (distributional) assumptions yields useful estimators (see below). Obtaining *S* from (1) is also referred to as the Horvitz-Thompson point estimate for zero-truncated distributions (Horvitz and Thompson, 1952).

#### Coverage estimates

The denominator in (1), 1 − *p*_0_ or 1 −ℙ[*X* = 0] = ℙ[*X* ≥ 1], reflects the proportion of observed species and is also called sample *coverage C*. Some of the estimators directly or indirectly estimate *C* as Ĉ and then Ŝ = *S*_obs_*/*Ĉ.

#### Distributional assumption: Poisson

As mentioned above, one may assume that 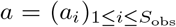 follows a certain type of probability distribution with probabilities *p*_*k*_ ≈ *f*_*k*_*/S* for *k* ≥ 0. Such an assumption cannot be justified in general, but may hold for certain types of datasets, and it simplifies the estimation problem. A popular distributional assumption for each *a*_*i*_ is the Poisson distribution, which models the random number *X* of successes when many attempts (*n* → ∞) are made, each with a very small success probability (*p* → 0), such that their product *λ* := *pn >* 0, corresponding to the expected number of successes, is a positive constant. Then, the Poisson distribution specifies that *p*_*k*_ =ℙ[*X* = *k*] = e^−*λ*^ · *λ*^*k*^*/k*!. The Poisson assumption can be exploited in different ways.

First, the Poisson distribution specifies that *p*_0_ = *p*_1_ = e^−*λ*^, so we can assume that *f*_0_ ≈ *f*_1_ and simply estimate Ŝ = *S*_obs_ + *f*_1_. This estimator can also be derived in a non-parametric way as a Jackknife estimator (see below).

Alternatively, under the Poisson assumption, ℙ[*X* = 1]*/*ℙ[*X* = 0] = *λ* can be estimated by *f*_1_*/f*_0_, and ?ℙ[*X* = 2]*/*?ℙ[*X* = 1] = *λ/*2 can be estimated by *f*_2_*/f*_1_. It follows that *f*_1_*/f*_0_ ≈ 2 *f*_2_*/f*_1_, or 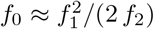, which is essentially the Chao 1 estimator (see the following section). Note that this estimator only uses *f*_1_ and *f*_2_ and not the other information contained in the data.

Still under the Poisson assumption, the data can be more comprehensively used if we compute an maximum likelihood estimate of the parameter *λ* from the observed zero-truncated Poisson distribution and then use (1) to estimate Ŝ = *S*_obs_*/*(1 − e^−*λ*^). This is the Poisson (PO) estimator (details in the following section).

### 2.4 Estimators in Detail

#### 2.4.1 Population Estimators

##### Good-Turing estimator (GT)

The Good-Turing estimator is one of the earliest richness estimators. Assuming that a *random* sample is drawn from an *infinite* population with a *finite* number of species *S*, Good (1953) proposed estimates for the probabilities that a species is represented exactly *r* times without making further assumptions about the population frequency distribution. One of their main results is that the proportion of species represented in the sample (coverage) is approximately Ĉ = 1−*f*_1_*/n*, or equivalently, the probability that the next observed individual belongs to an unseen species is given by 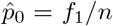. This result led to the common assumption that rare species, especially the number of singletons, contain most information about the number of missing species.

Good (1953) did not further comment on predicting the species richness given the probability to observe a new element. However, we may use the estimate 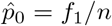 (or Ĉ = 1 − *f*_1_*/n*) together with (1) or the coverage estimate to obtain the GT estimator

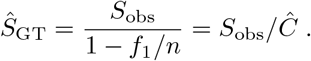

##### Jackknife estimators (Jack 1, Jack 2)

Burnham and Overton (1978) derived *non-parametric* estimators that are a linear combination of the species frequencies. The derivation is based on the following assumptions: the population is *closed*, the species detection rate is *constant* for each species but may vary between species and the capture events are all *independent*. It follows that the observed capture frequencies are a *random* variable following a *multinomial* distribution with unknown success probabilities (Burnham and Overton, 1979). Instead of assuming a parametric distribution for the success probabilities, Burnham and Overton (1978) derive a non-parametric estimator using the generalized Jackknife method. The first and second order Jackknife estimators are given by

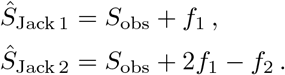

##### ACE estimator (ACE)

The abundance-based coverage estimator (ACE) is a modification of the GT estimator in the sense that it considers only *rare* species for the coverage estimator. The species are separated into rare and abundant groups based on a frequency cutoff *T*, i.e., species are rare if they are observed at most *T* times (Chao, 2006). The most common cutoff is *T* = 10, but results are sensitive to the choice of *T*.

We apply *Ŝ*_GT_, but instead of using the entire number *S*_obs_ of observed species, we only use the number *S*_rare_ of rare species and adjust *n* accordingly. The number of abundant species is simply counted as-is. We obtain

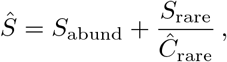

Where

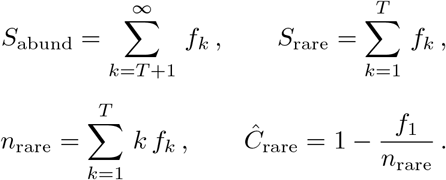

The above estimator is not the final ACE estimator because it assumes that all rare species are *homogeneous*, i.e., all species are assumed to have the same relative abundance. Since the homogeneity assumption may be violated, Chao and Lee (1992) proposed an adjusted estimator that accounts for the *heterogeneity* of rare elements. For a population with true relative species abundances (*q*_*i*_)_1≤*i*≤*S*_, the abundance distribution may be summarized by their mean 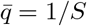 and coefficient of variation (CV), where the squared CV is defined as

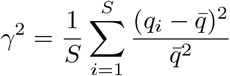

Based on the results by Good and Toulmin (1956), Chao and Lee (1992) estimate *γ*^2^ by

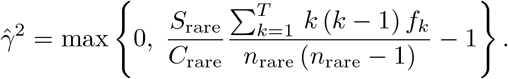

With this estimate of *γ*^2^, Chao and Lee (1992) obtained

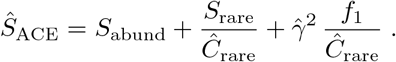

We point out again that the estimate is sensitive to the rareness abundance threshold *T*.

##### Poisson estimator (PO)

The PO estimator has already been briefly introduced above: We assume that each 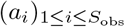 is drawn from a Poisson distribution with an unknown parameter *λ >* 0. We estimate *λ* from the zero-truncated distribution as 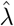 and then use (1) to estimate

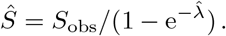

We now provide details on the estimation of *λ* using the maximum likelihood approach on the zero-truncated Poisson distribution, where for *k* ≥ 1,

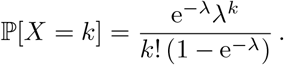

For a sample of size *n* and observed species abundances 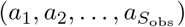.the likelihood function is given by

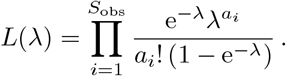

It follows that the MLE must satisfy

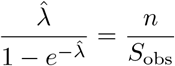

which can be solved numerically for 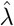 (van der Heijden et al., 2003).

Instead of considering all species to estimate 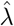, the sample may first be restricted to rare species with an abundance bounded by a user-defined threshold *T* (default *T* = 10), with the same impliciations as for the ACE estimator.

##### Chao 1 estimator (Chao 1, Chao 1-BC)

As introduced in Section 2.3, Chao (1984) proposed a *nonparametric* estimator that can be derived under the assumption that the observed species counts follow a Poisson distribution with equal detection rate *λ*. The estimator is given by Chao (1984) proved that the estimator yields a lower bound of the true richness for *n* → ∞ under both multinomial and Poisson models. The lower bound can be derived based on the monotonicity of the ratio of consecutive probabilities (Chao, 1987; Böhning et al., 2013).

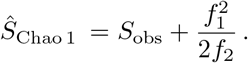

Since the Chao 1 estimator is undefined for *f*_2_ = 0, it has been replaced in later work (Chao, 2006) by a bias-corrected version, given by

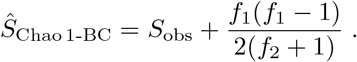

Its derivation requires additional assumptions, including species homogeneity, which often do not hold in practice (Böhning et al., 2019).

A hybrid form is to use *Ŝ*_Chao 1_ for *f*_2_ *>* 0 and the bias-corrected term *S*_obs_ + *f*_1_(*f*_1_ − 1)*/*2 if *f*_2_ = 0.

##### Gamma-Poisson mixture estimator (GPM)

Under the assumption that the detection rate varies between species, the Poisson parameter *λ* is itself a random variable. If we assume that the species-specific *λ*_*i*_ are drawn from a Gamma distribution, then the observed species counts follow a Gamma-Poisson mixture distribution, which is a common assumption of many richness estimators (Chao and Bunge, 2002; Lanumteang and Böhning, 2011; Chiu, 2023).

The marginal distribution of a Gamma-Poisson mixture model with parameters *α* and *β* is given for *x* = 0, 1, 2, … by

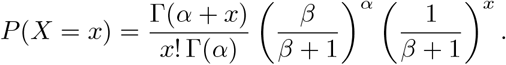

Under the zero-truncated Gamma-Poisson mixture model, the probability *p*_*k*_ to observe a species exactly *k* times is given by *P* (*X* = *k*)*/*(1 − *P* (*X* = 0)).

As introduced in Section 2.3, the Horvitz-Thompson richness estimator is given by

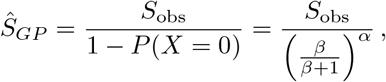

which requires the estimation of *α* and *β*.

The observed frequencies *f*_*k*_ follow a multinomial distribution with total sum *S*_obs_ and probabilities (*p*_*k*_)_*k*≥1_. Hence, we may solve for *α* and *β* by maximizing the likelihood function

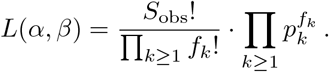

For detailed information about deriving the MLE for a Gamma-Poisson mixture distribution, we refer the reader to the paper by Chiu (2023).

##### Chao and Bunge estimator (CB)

Chao and Bunge (2002) proposed an estimator that has a *non-parametric* form, but the optimality criteria hold under a GammaPoisson mixture model.

For a sample with observed richness *S*_*obs*_ and number of individuals for each species in the sample given by 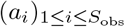, the Chao and Bunge estimator is given by

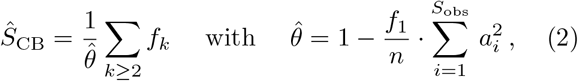

which is based on a consistent estimator for the expected value of *f*_0_ under a Gamma-Poisson mixture model.

##### Lanumteang and Böhning estimator (LB)

Lanumteang and Böhning (2011) derived a species richness estimator by computing a Taylor expansion over the log-ratios log(*j p*_*j*_*/p*_*j*−1_), where *p*_*j*_ has a Gamma-Poisson mixture distribution. Solving the equations in *j* = 1, 2 for *f*_0_ allows to derive an estimate for *f*_0_ that is *non-parametric* in form; given by

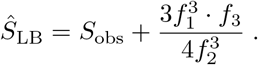

##### Chiu estimator (Chiu)

Chiu (2023) proposed a moment estimator for a Gamma-Poisson mixture model that estimates the parameters based on the expected values for the number of unseen species, singletons, doubletons and tripletons. The derived point estimate to predict species richness is given by

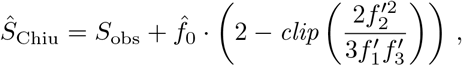

where 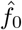 is estimated using the Chao 1 estimator, 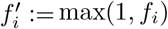, and *clip*(*A*) := min(max(1*/*2, *A*), 1), i.e., *A* clipped to the interval [1*/*2, 1].

##### Objective Bayesian estimator (OB-PO, OB-NB, OB-G, OB-MG)

Under the assumption that the species abundances follow a certain parametric probability distribution, the parameters may alternatively be estimated using Bayesian statistics instead of maximum likelihood approaches, for example using Bayesian estimators that place an objective prior on the number of species and their frequency distributions.

Barger and Bunge (2010) suggest to use reference priors, which maximize the expected entropy. In this review, we evaluated the Bayesian estimators for a Poisson (OB-PO), negative binomial (OB-NB), geometric (OB-G) and mixed geometric (OB-MG) distribution.

##### Recon

To estimate diversity of B- and T-cell repertoires, Kaplinsky and Arnaout (2016) developed Recon, a maximum likelihood approach that makes no parametric assumption about the frequency distribution.

The estimator is calculated using an expectation-maximization approach that adds in each iteration new parameters until further parameters would lead to overfitting. The algorithm starts from a uniform frequency distribution, i.e., all species have the same number of individuals in the population. In each iteration, a new species frequency is added and the respective species counts and relative frequencies are fitting by maximum likelihood. Apart from species richness, the predicted frequency distribution may be used to estimate other diversity measures, such as entropy, the Gini-Simpson index or the Hill numbers.

##### Valiant

Valiant and Valiant (2017) developed a linear program that estimates the shape of the unobserved portion of the frequency distribution. They show that their approach yields accurate results for various natural distributions if the sample size is at least in *O* (*S/* log *S*) for a population with *S* distinct elements.

The algorithm is a combination of two linear programs. The first linear program searches a histogram whose expectation is closest to the observed frequency distribution. The second linear program optimizes the objective of finding a histogram that has minimal support size under the constraint that the new histogram has a similar distance to the observed frequency distribution as the one obtained by the first linear program. The coefficients to calculate the expected frequencies are computed using Poisson probabilities.

##### Breakaway

Willis and Bunge (2015) estimate species richness using a heteroscedastic, correlated nonlinear regression model to fit ratios of consecutive frequencies. By fitting a rational to the ratios of the form *f*_*j*+1_*/f*_*j*_ as a function of *j*, the estimate for the number of unseen elements *f*_0_ is given by projecting the fitted function to 0.

Since a robust estimate for the number of missing species requires an accurate number of singletons, Willis (2016) enhanced the previous approach by predicting both the number of unobserved elements and the number of singletons, called Breakaway-f1.

#### 2.4.2 Upsampling Estimators

The estimators in the previous section 2.4.1 attempt to estimate the species richness *S* of the full population without knowing its size *N*, even allowing infinite *N*, but assume that *S* is finite. In contrast, the estimators in this section are given the population size *N* as additional input, and therefore do an *upsampling* estimate (from observed richness *S*_obs_ and (*f*_*k*_)_*k*≥1_ with *n* individuals to the unknown *S* with *N* individuals). The difficulty of the upsampling problem increases with the ratio *N/n*.

#### Good-Toulmin estimator (EF-GT, PO-GT)

To estimate how many unseen species may be expected in a next sample, Good and Toulmin (1956) proposed an estimate that is based on the assumption that the capture rates follow a Poisson distribution. Since the probability of observing a new species is given by the probability that a species was not seen in the first but was seen in the second sample, the expected number *E* of new species one would see in a second sample *of same size* is given for a population with *S* species by

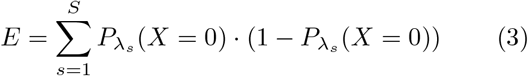

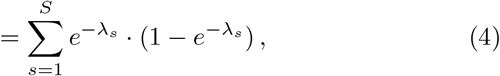

where 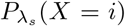 denotes the probability to observe exactly *i* individuals from species *s* under a Poisson distribution with parameter *λ*_*s*_. The series expansion of the second term is given by

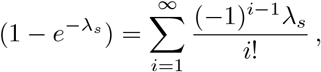

which may be used to rewrite (4) as

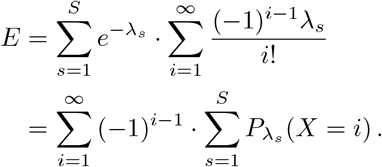

Good and Toulmin (1956) approximate the sum 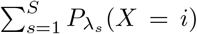 by the observed *f*_*i*_ and generalize the procedure to general sizes *m* of the second sample.

They obtain an estimate for the number of unseen elements Û in a new random sample of size *m* from sample of size *n:*

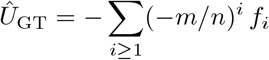

with ratio *t* = *m/n*. Note that *t* = 1, as derived above, corresponds to upsampling by a factor of 2, where the second sample has the same size as the first, and *t* = 99 corresponds to 100x upsampling. They showed that the above formula is a nearly unbiased *non-parametric* estimator of *U* for all *t* ≤ 1, but the convergence of the series is not guaranteed for *t >* 1.

One possibility to achieve convergence for *t >* 1 is to perform so-called probabilistic smoothing, yielding new estimators of the form

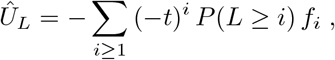

where the discrete random variable *L* can follow an arbitrary discrete distribution.

Efron and Thisted (1976) proposed Binomial smoothing (GT-ET), such that

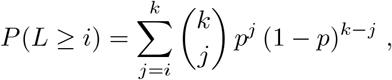

With 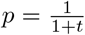, and *k* being a tuning parameter. We use 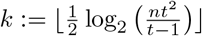,which has been shown by Orlitsky et al. (2016) to lead to the best convergence rate.

As an alternative, Poisson smoothing (GT-PO) uses

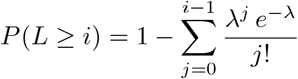

with 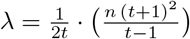.(Orlitsky et al., 2016).

The resulting estimates Û are in fact estimates of *f*_0_; thus the richness estimator is *Ŝ* = *S*_obs_ + Û. This also holds for the following estimators, which use more complex procedures to estimate the number of unseen species.

##### PreSeq

To approximate the molecular complexity of sequencing libraries, Daley and Smith (2013) introduced the idea of using rational function approximation to increase the radius of convergence of the Good-Toulmin power series. Rational function approximation increases the radius of convergence for divergent series, in particular for alternating power series such as the Good-Toulmin power series, which allows to predict the true richness for samples that are several orders of magnitudes larger than the original sample.

In addition, they first apply the Euler transform proposed by Good (1953); yielding the power series

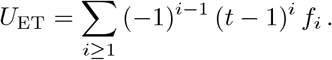

By transforming the variable *t* of the power series, Pre-Seq considers a larger class for the rational function approximation, but under the constraint that the first coefficients are equal to the coefficients of the original power series, hence trusting the original series more in the neighborhood around *t* = 1 (Daley and Smith, 2013).

##### Pitman/Ewens sampling formula (PSF)

The Pitman sampling distribution, a two-parametric generalization of the Ewens sampling formula, is a common sampling model that assumes an infinite sampling universe (Pitman, 1995). The urn representation of the Pitmam sampling formula is given by the Hoppe urn model. The formula calculates the probability of observing an integer partition of *n*, where a partition is assumed to be random and exchangeable. The set of valid integer partitions is given by 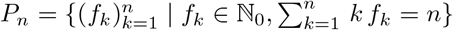 .The probability of a partition *f* under a Pitman sampling model with parameters *α* ∈]0, 1] and *θ* ∈ [−*α*, ∞] is defined as

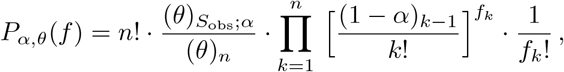

where (Pitman, 1995)

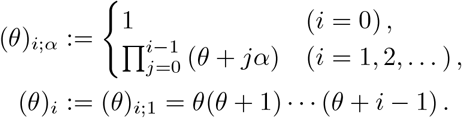

To estimate species richness, the parameters *α* and *θ* of the Pitman sampling distribution can be estimated using maximum likelihood estimation. Given *α* and *θ*, the expected number of additional elements Û_PS_ in a next sample of size *m* is given by (Tauber and Haeseler, 2013)

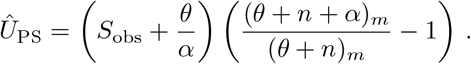

##### DivE

Rarefaction curves show the number of distinct elements as a function of the sample size and are a common method in diversity estimation to analyze species richness via nested subsampling. Traditional curve fitting approaches fit a parametric asymptotic function, such as a negative exponential or logistic function, to the rarefaction curve (Colwell et al., 1997). The species richness is then given by the asymptote of the function. Laydon et al. (2014) extended this idea by fitting a list of 58 mathematical function classes to the rarefaction curves and accumulating the results of the five best fitting classes. Instead of computing the asymptote, the predicted species richness is given by extrapolating each function to the desired sample size. Since testing all mathematical functions is very compute intensive, we restricted our evaluation to previously suggested functions: the logistic, negative exponential, logarithmic, hyperbolic and Hill function families.

##### RichnEst

Schröder and Rahmann (2015) developed a linear program (LP) to estimate the molecular complexity or duplication rate of sequencing experiments from a small sample. The linear program searches for plausible frequency vectors of the whole population by minimizing the distance between the expected frequency vector and the observed frequency vector of the sample.

Assuming that the sample is drawn randomly, the probability that a species with *K* individuals in the complete population of size *N* is observed exactly *k* times in a sample of size *n* follows a hypergeometric distribution, which allows to compute the expected frequency vector, assuming a population frequency vector (Schröder and Rahmann, 2015). The LP formulation is used to invert this forward downsampling process.

## Evaluation

### Data Simulation

The richness estimators were evaluated on simulated data with nine different species compositions (Fig. 2). We considered samples containing 1%, 3%, 5%, as well as 10%, 20%, …, 90% of the population. For each test case, a sample was drawn 20 times, resulting in a total of 2160 test cases.

**Figure 2.**
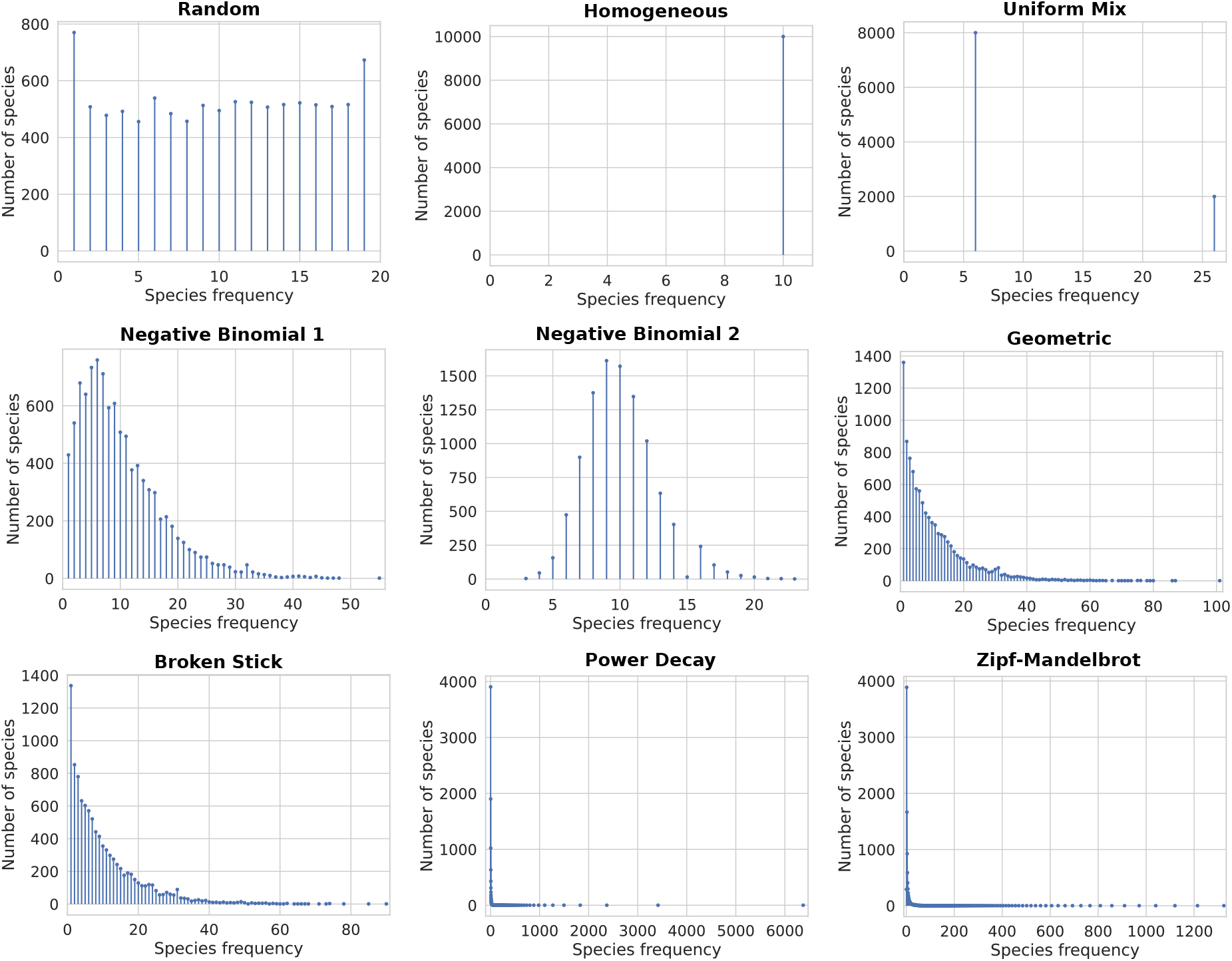
Simulated data. Each plot shows the species frequency distribution of a population with 10^5^ individuals and 10^4^ distinct species under different population models.

For each species composition, the relative abundances for a population with *N* = 10^5^ individuals and *S* = 10^4^ distinct species are given (*p*_1_, *p*_2_, …, *p*_*S*_) = (*cA*_1_, *cA*_2_, …, *cA*_*S*_), such that 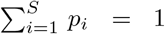 and 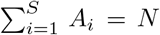, where the *A*_*i*_ are the absolute population abundances and *c* = 1*/N* (Chiu, 2023). Below, we specify *p*_*i*_ or *A*_*i*_ of different population composition models. Data simulation and evaluation was automated using Snakemake (Mölder et al., 2021).

#### Random model

For *i* = 1, …, *S*, we draw *p*_*i*_ from the uniform distribution on [0, 1[, normalize such that 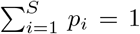 and multiply by the population size *N* to obtain the abundances *a*_*i*_, with an average of 10. Values *a*_*i*_ = 0 are increased to *a*_*i*_ = 1. To compensate, a different species *i* with *a*_*i*_ = 20 is reduced to *a*_*i*_ = 19.

#### Homogeneous model

For *i* = 1, …, *S*, we set *p*_*i*_ = 1*/S*. Thus, for *N* = 10^5^ and *S* = 10^4^, each species has exactly 10 individuals.

#### Uniform mixture model

For one fifth of the species, *i* = 1, …, *S/*5, we set *p*_*i*_ = 5*/*(2*S*), and for the remaining 4/5 of the species, *i* = *S/*5 + 1, …, *S*, we set *p*_*i*_ = 5*/*(8*S*) (assuming that *S* is a multiple of 5).

#### Negative binomial models

We use two different models. For *i* = 1, …, *S*, we set 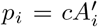,where 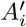 is a random sample drawn from a negative binomial distribution either with parameters *r* = 2 and *p* = 0.02 (model 1), or with *r* = 20 and *p* = 0.2 (model 2).

#### Geometric model

For *i* = 1, …, *S*, we set 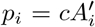,where 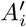 is a random sample drawn from a geometric distribution with parameter *p* = 1*/S*.

#### Power decay model

For *i* = 1, …, *S*, we set *p*_*i*_ = *c/i*^0.9^, where *c* is the proper normalization constant.

#### Zipf-Mandelbrot model

For *i* = 1, …, *S*, we set *p*_*i*_ = *c/*(*i* + 10), where *c* is the proper normalization constant.

#### Broken stick model

For *i* = 1, …, *S*, we set 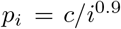,where *A*^*′*^ is a random sample drawn from a standard exponential distribution.

### Real Datasets

We applied the tools to publicly available immune repertoire and mircobiome sequencing datasets.

We used the repertoire sequencing data provided by Shugay et al. (2015, VDJTools Examples). Targeted repertoire sequencing refers to the sequencing of V(D)J genes which code for antibodies and T-cell receptors. The clonotype of each cell can be inferred based on the sequencing information, allowing a high level analysis of immune repertoire diversity. We applied the tools to 58 available samples, for which we created 10 subsamples for each of 6 subsampling rates of 1%, 3%, 5%, 10%, 20% and 30%, resulting in 3480 test cases.

To estimate microbiome diversity, we applied the tools to 20 metagenomic datasets from Durazzi et al. (2021, MG-Rast database), which contain the abundance of bacterial strains in the chicken gut. For each dataset, we created the same 10 × 6 subsample types as for the immune repertoires, resulting in 1200 test cases.

## Results

The tools require as input either the observed abundance vector 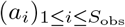, i.e., the number of times each species was observed in the sample, or the frequency vector (*f*_*k*_)_*k*≥1_, i.e., the number of species occurring exactly *k* times in the sample, and output a point estimate and sometimes a confidence interval of the population species richness. Since not all tools compute a confidence interval, we measure the accuracy based on the point estimates.

On simulated data, we evaluate the tools with respect to (1) proportion of crashes, (2) proportion of outliers, (3) point estimation accuracy and (4) computational resource requirements. Next, we evaluate the point estimation accuracy for the V(D)J and microbiome subsamples.

### Proportion of Unsolved Problems

Figure 3A shows the proportion of unsolved test problems for each tool. A test problem is unsolved if the tool either crashed or failed to converge to a solution. Most of the tools are able to compute a point estimate for all problems, except for the objective Bayesian estimators and smoothed Good-Toulmin estimators which sometimes fail to converge. In addition, the Good-Turing and Lanumteang-Böhnung estimator fail if *f*_1_ or *f*_2_ are zero, respectively. A similar problem holds for the Chao-Bunge estimator if 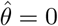 in (2). For detailed information about the proportion of unsolved problem per subsampling rate and population, see Supplementary Figure S.2.

**Figure 3.**
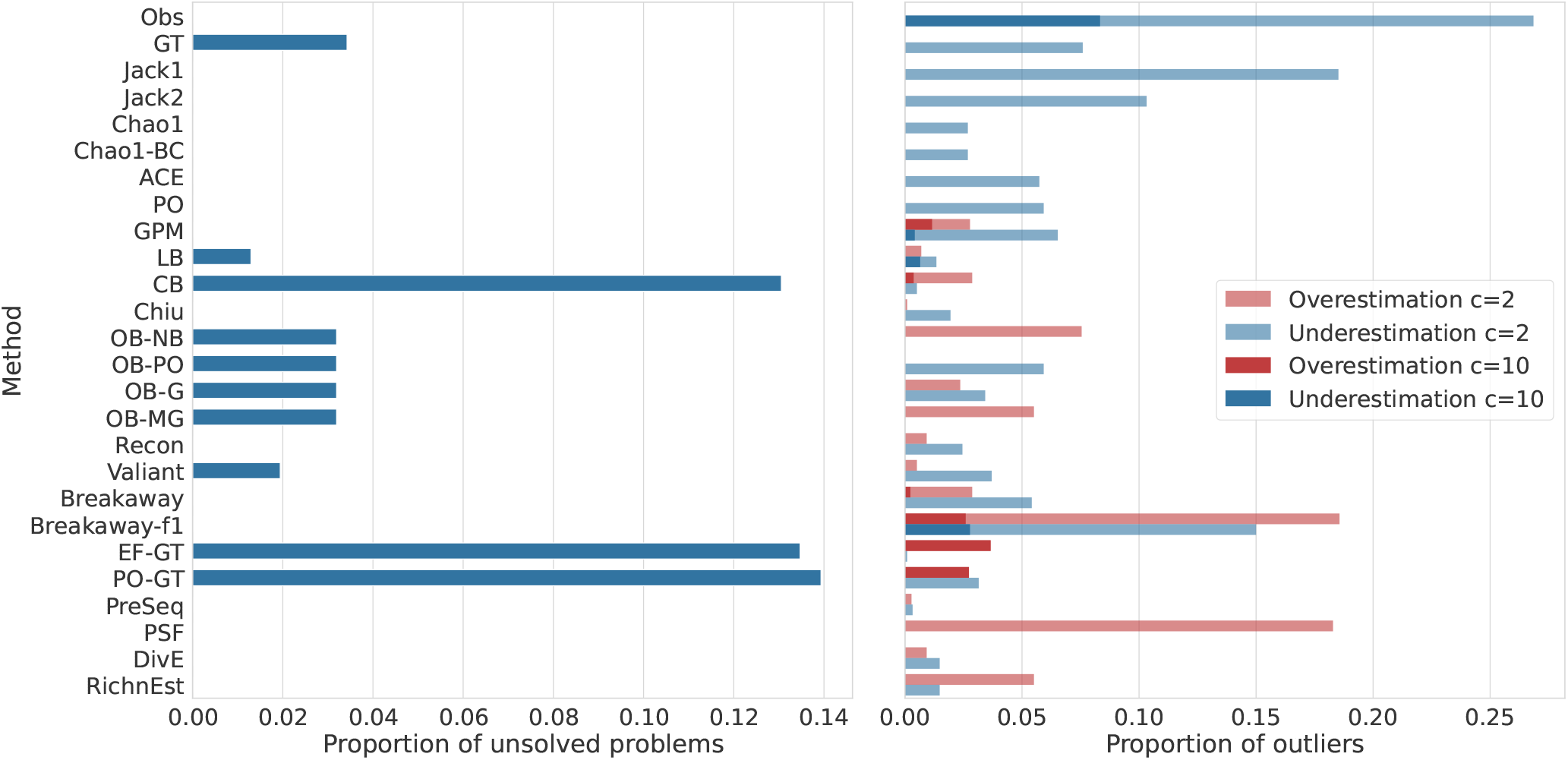
**A** For each tool, the proportion of unsolved problems is shown. Lower is better; zero is desirable. **B** Proportion of outliers for each tool. For constant *c* ∈ {2, 10}, each point estimate that is smaller than *S/c* or larger than *cS* is assumed to be an outlier that heavily under- or overestimates the true population richness, respectively. Lower is better.

### Proportion of Outliers

We compared the number of outliers per tool, where we consider an estimate to be an outlier if *Ŝ < S/c* or *Ŝ > cS*, for a constant *c >* 0. Figure 3B shows the proportion of outliers for *c* = 10 and *c* = 2.

As expected, the observed species richness (Obs) often strongly underestimates the true richness. For several test cases, Breakaway-f1 and the maximumlikelihood based Poisson and Gamma-Poisson estimators strongly over- and underestimate the true richness. In addition, the Pitman sampling formula and the smoothed Good-Toulmin estimators strongly overestimate the true richness for small sampling rates. For the Good-Toulmin estimators this may be caused by divergence of the power series. In general, simple estimators, such as Chao1, usually underestimate the true richness, while more complex estimators both under- and overestimate the true richness (see Figure 3B). Moreover, all tools have most outliers for small sampling rates (1% to 5%). For larger sampling rates, populations with a Power Decay and Zipf-Mandelbrot distribution are most challenging (see Supplementary Figure S.3).

### Estimation accuracy

Figure 4 gives a more detailed overview of the estimation accuracy. Each panel corresponds to a subsampling rate.

**Figure 4.**
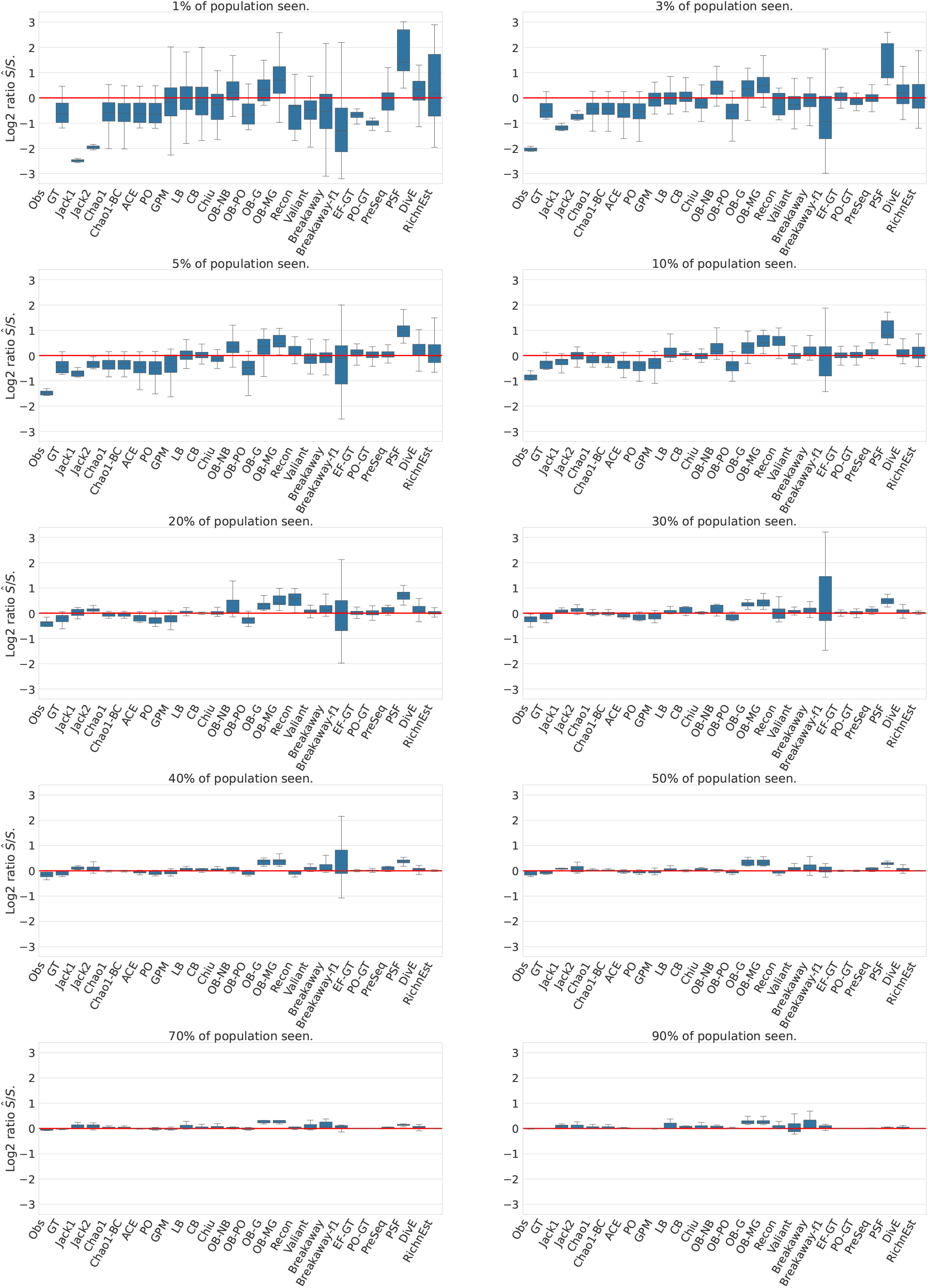
Boxplots of estimation accuracy on simulated data. Shown are log_2_ ratios between the estimated species richness and the true species richness for different subsampling rates (panels), combined over all population models. If the prediction is correct, log_2_ *Ŝ/S* = 0 (red horizontal line). If log_2_ *Ŝ/S >* 0, the estimator overestimates the true richness and if log_2_ *Ŝ/S <* 0, the estimator underestimates the true richness. Failures and outliers (using *c* = 10; see Fig. 3) are not included in this data.

**Figure 5.**
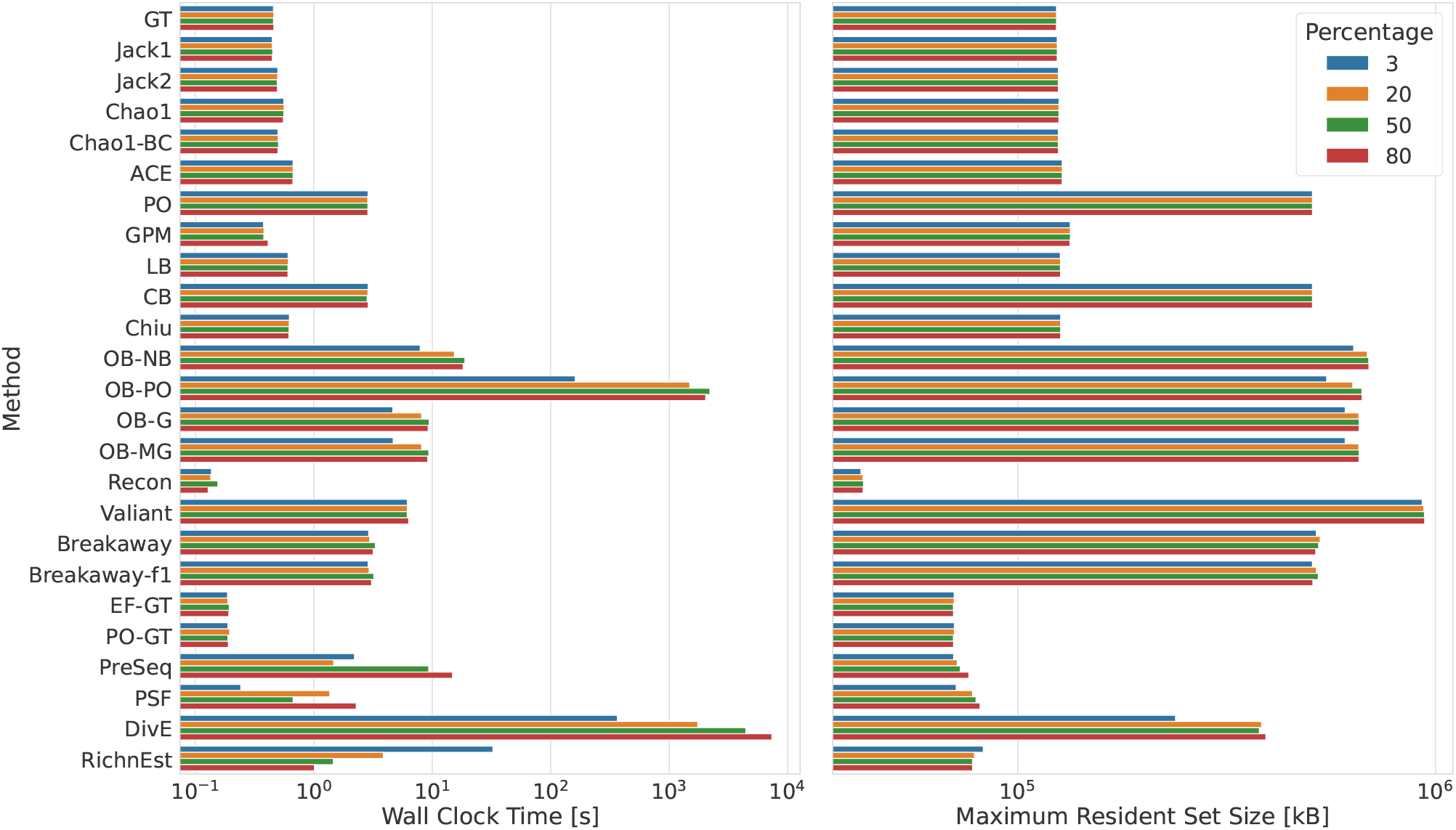
Computational resource requirements: **A** Wall clock time in seconds; **B** Memory usage in kilobytes. The benchmarks for a sample containing 3%, 20%, 50% and 80% of the total population were averaged over three runs for populations with a negative binomial (*r* = 20 and *p* = 0.2) and Zipf-Mandelbrot frequency distribution on a AMD Ryzen 9 5950X 16-Core processor with a maximum CPU clock speed of 5.1 GHz.

Each box plot shows the deviation of the point estimate from the true richness across all population models for one tool. Individual plots for each population model are provided in supplementary figures.

In general, the estimation problem is more challenging when only a small portion of the population has been observed, resulting in an underestimation of the true richness by most tools, such as the Chao 1 estimator, the ACE estimator or Valiant. When the sample contains more than 50% of the population, methods that do not require an upsampling factor sometimes overestimate the true richness.

The non-parametric and Gamma-Poisson mixture estimators give accurate results for populations with a negative binomial and homogeneous frequency distribution, but tend to underestimate the true richness for populations under a geometric, Zipf-Mandelbrot or power decay model (Supplementary Figures S.4-S.12), with the Chao 1, ACE and Chiu estimators being among the most accurate methods.

Breakaway and Breakaway-f1 both over- and underestimate the true richness and have large estimation intervals for repeated samples (Supplementary Figure S.7). In addition, Recon and the Bayesian Geometric and Mixed Geometric estimators tend to overestimate the true richness. Even for a population with a geometric frequency distribution, the objective Bayesian geometric and mixed geometric estimators consistently overestimate the true richness (Suppl. Fig. S.9).

Upsampling methods, that require the total population size as an additional parameter, show increasing accuracy with increasing sample size. For example, comparing the linear based programming tools RichnEst (upsampling estimator) and Valiant (population estimator) shows that RichnEst yields more accurate results than Valiant if the subsampling rate is higher than 40%, while Valiant tends to overestimate the true richness. However, if the sample contains less than 10%, Valiant outperforms RichnEst. In particular for populations with many rare species, such as the geometric, power decay and Zipf-Mandelbrot models, RichnEst yields inaccurate results for small samples.

The smoothed Good-Toulmin estimators provide accurate results for many of the evaluated problems, even for small samples. However, they can suffer from convergence problems, e.g., many of the problems could not be solved by the smoothed Good-Toulmin estimators if less than 30% of a population with a power decay frequency distribution was observed (see Supplementary Figure S.2). PreSeq’s approach of using rational function approximation successfully increases the convergence ratio of the power series. PreSeq is able to solve all problems and is among the best performing tools for populations with a power decay or Zipf-Mandelbrot distribution (Supplementary Figure S.12). However, is outperformed by non-parametric estimators for a population with a negative binomial frequency distribution (Supplementary Figure S.6). The performance of DivE is similar to PreSeq showing an increasing accuracy for higher subsampling rates.

Although the Pitman sampling formula also requires an upsampling factor, it is less accurate than most population and upsampling estimators, i.e., the true richness is often overestimated (see Figure 4).

### Computational Requirements

Running times range from under one second for tools such as the Good-Turing, Jackknife, Chao 1 and Gamma-Poisson mixture model, to an hour for the Objective Bayes Poisson estimator and several hours for DivE. Although we evaluated DivE with only 5 different mathematical function families, the running time was already significantly higher compared to the other approaches. In addition, DivE’s time requirements strongly increase with increasing sample size, which makes it impractical for large datasets.

The maximum memory requirements are under 1 GB for all tools and evaluated sample sizes and largely independent of sample size.

### Estimating V(D)J Diversity

Methods which could not solve all problems (Good-Turing, Chao-Bunge, Objective Bayesian, smoothed Good-Toulmin), had extreme outliers (Breakaway-f1, Pitman sampling formula), performed less accurate for most problems (Jackknife 1 and 2, Poisson model) or have impractical computation times on large data (DivE, e.g., 120h for one V(D)J dataset and a subsampling rate of 0.3) are not considered in the subsequent evaluation.

Figure 6 shows the deviation of the predicted species richness from the observed species richness in the complete V(D)J sequencing dataset for different subsampling rates. The boxplot labelled “Obs” shows the number of observed clonotypes in the subsample; it always underestimates the true richness. For the population estimators, the observed richness of the complete dataset imposes a lower bound on the true V(D)J richness, but the true repertoire richness may be considerably higher considering that only a small subset of immune cells was sequenced. By contrast, the upsampling estimators PreSeq and RichnEst should be close to the observed richness of the complete dataset.

**Figure 6.**
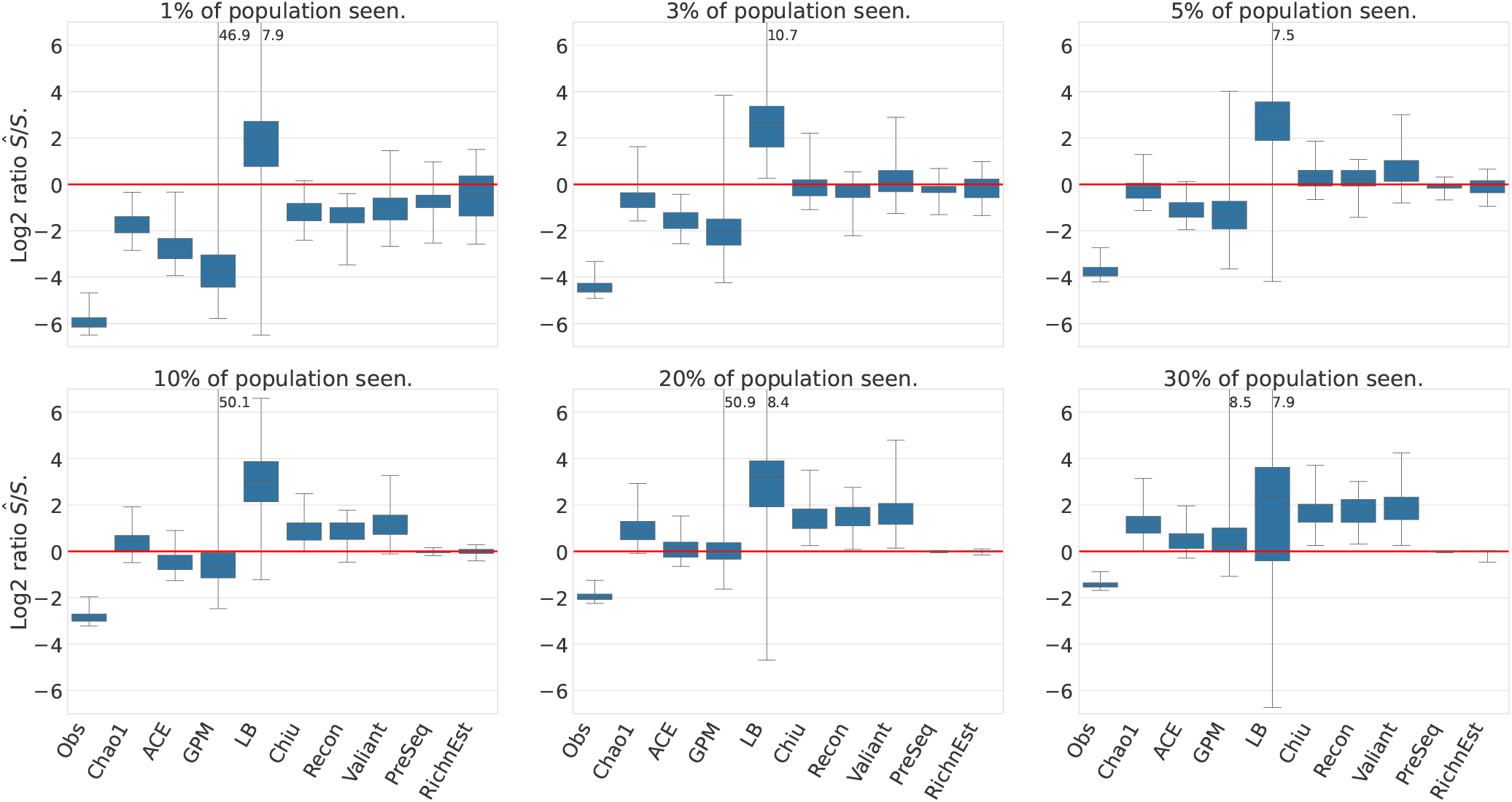
Estimation of V(D)J diversity from repertoire sequencing data. Boxplots displaying the distribution of the log_2_ ratios between the predicted species richness and the species richness observed in the complete dataset. Numbers at the cut whiskers show the maximum deviation of the method.

At low subsampling rates, Chao 1, ACE and the Gamma-Poisson mixture model strongly underestimate the true richness, which is consistent with the previous results that these estimators tend to underestimate the true richness for compositions with many rare elements. The predictions by Chiu, Recon and Valiant are close to the true richness of the full sample if 3% of the population has been observed, and they predict a higher estimate for higher subsampling rates. Since the true richness of the immune repertoire is unknown (our 100% is in fact also only a subsample of unknown proportion), the accuracy cannot be validated, but the almost linear rarefaction curves indicate that the total VDJ diversity is substantially larger than the full sample richness (see Supplementary Figure S.13). This equally holds for the Lanumtang-Böhnung population estimator, which is the only method that predicts an extremely higher richness for all subsampling rates.

The point estimates of RichnEst vary more compared to PreSeq. However, for a subsampling rate of 3% and 5% the predictions of RichnEst are on average closer to the full richness. Both tools are close to the full richness, with an increasing accuracy for larger subsamples.

### Estimating Microbiome Diversity

Durazzi et al. (2021)’s metagenomics data allows us to evaluate the performance of richness estimation on data where the true population richness is close to the sample richness (see rarefaction curves in Supplementary Figure S.13). Figure 7 shows that all methods underestimate the species richness for subsampling rates below 10%. For a subsampling rate of 10% and 20% only the Lanumtang-Böhnung estimator and Valiant predict a higher population richness, and Recon strongly underestimates the species richness. All other methods have a similar performance, with PreSeq yielding the most accurate results.

**Figure 7.**
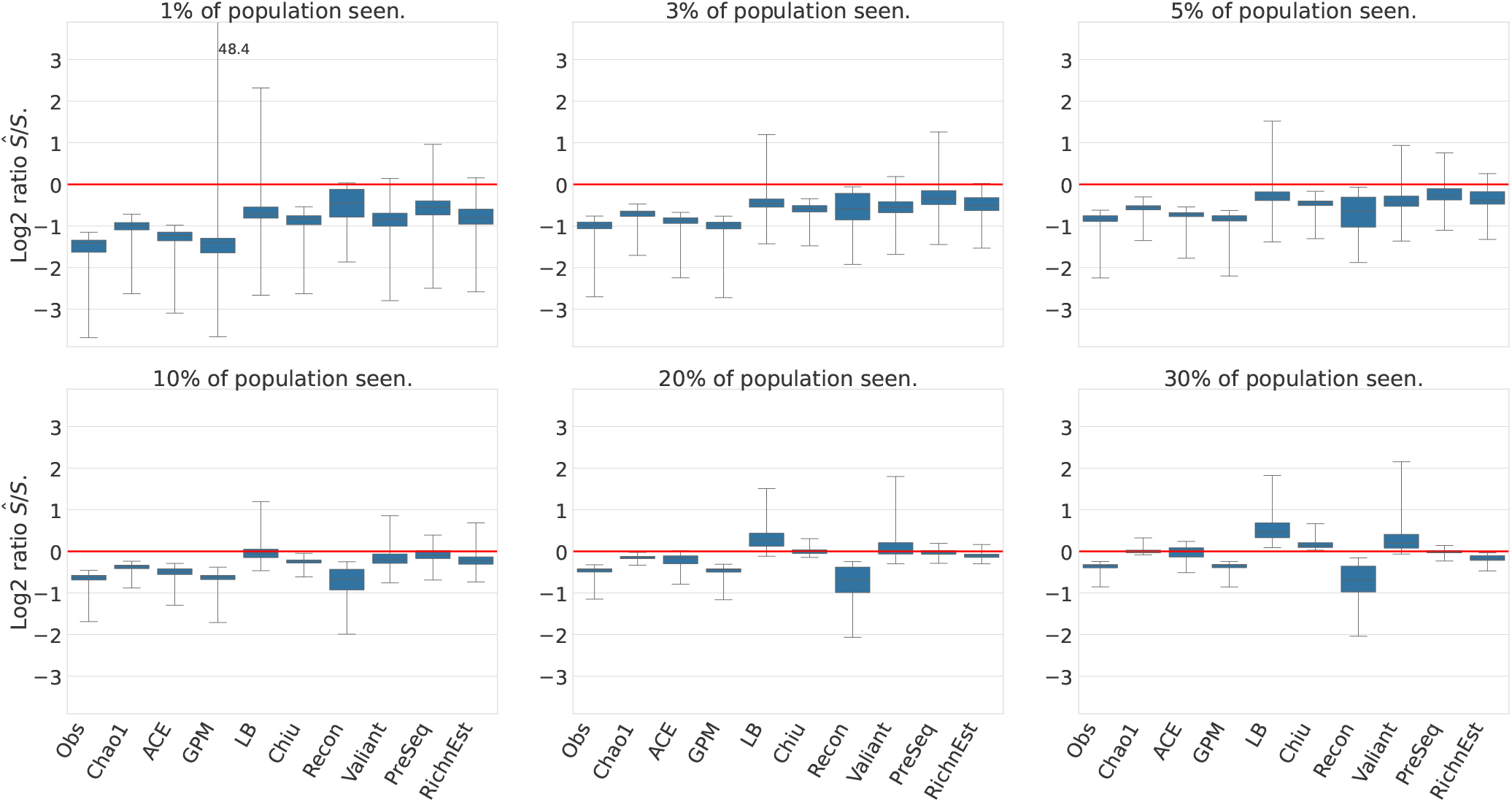
Estimation of gut microbiome species richness from whole shotgun metagenomic sequencing data. Boxplots display the distribution of the log_2_ ratios between estimated species richness and species richness in the full dataset. Numbers at the cut whiskers show the maximum deviation of the method.

## Conclusions

The increasing applications of richness estimation, for instance estimating the diversity of immune repertoires, make the accurate estimation of species richness from a small sample an important research topic. Although a variety of richness estimators already exist, new approaches that are either specific or generally applicable to many different scenarios, are still being developed. We presented an overview of existing richness estimators and evaluated them on a wide range of populations with different species compositions.

Richness estimators are classified into population and upsampling estimators, depending on whether they require the size *N* of a future sample as an additional parameter. Among the population estimators, the estimators that are non-parametric in form, like the Chao 1, ACE and Chiu estimator, provided the most accurate point estimates for a variety of species composition, but underestimated the true species richness if the population was either very heterogeneous are only a small portion of the sample had been observed. The estimates of more complex methods, like Breakaway or the Objective Bayesian estimators, often had a high variation for repeated subsamples from the same population and both strongly over- and underestimated the true richness.

Even tough upsampling estimators have the benefit of knowing the total population size, they did not consistently perform better than the population estimators. For example, the Pitman sampling formula yielded inaccurate results for many problems, even if the sample contained more than half of the population. However, PreSeq and RichnEst yielded accurate results for most of the problems and often outperformed the other approaches. RichnEst provided accurate results when more than 10% of the population was contained in the sample, but suffered from inaccuracies when the sample was too small; in these cases PreSeq performed better than RichnEst.

In conclusion, our evaluation shows that different estimators performed best for different species compositions. Overall, PreSeq, RichnEst and the Chiu estimator were most accurate among different species compositions.

## Author contributions

The evaluation was done by J.S. under the supervision of S.R.. J.S and S.R. contributed to writing and revising the manuscript.

## Ethics declaration

None declared.

## Data availability

The code for simulating and downloading the data is part of a Snakemake workflow available at https://gitlab.com/rahmannlab/speciesrichness.

## Supplementary Figures

**Figure S.1:**
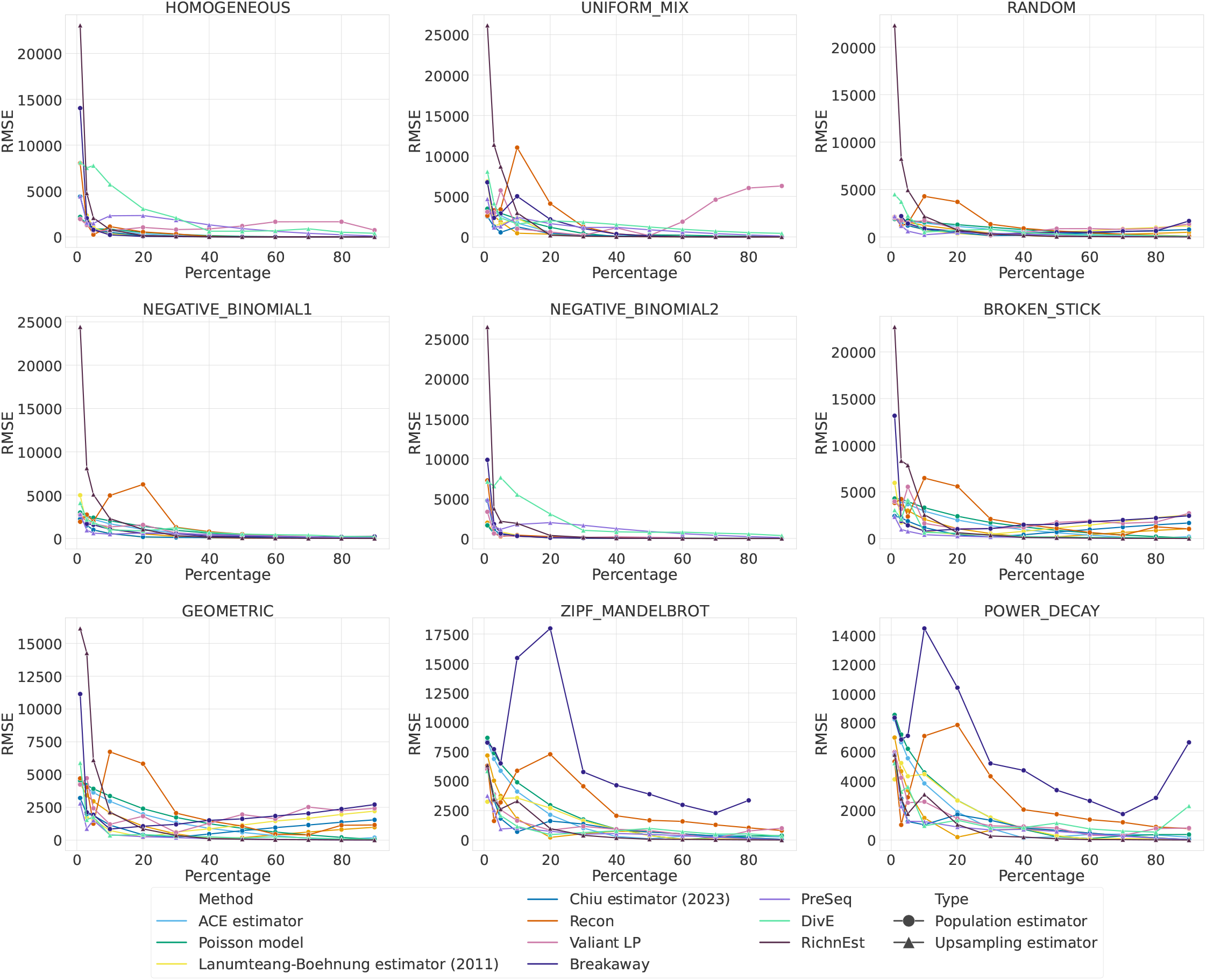
Root mean square error (RMSE) changes with increasing sample size across population models. The RMSE is defined for *k* samples with true species richness *S*_1_, …, *S*_*k*_ and estimated species richness *Ŝ* _*1*_, …, *Ŝ*_*k*_ as 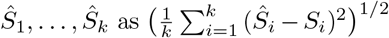. Estimators or tools were not considered if they could not solve most problems (Good-Turing, Chao-Bunge, Objective Bayesian, smoothed Good-Toulmin), had many outliers (observed richness, Gamma-Poisson mixture model, Breakaway-nof1, UnseenEst) or performed badly for most problems (Jackknife 1 and 2, Pitman sampling formula).

**Figure S.2:**
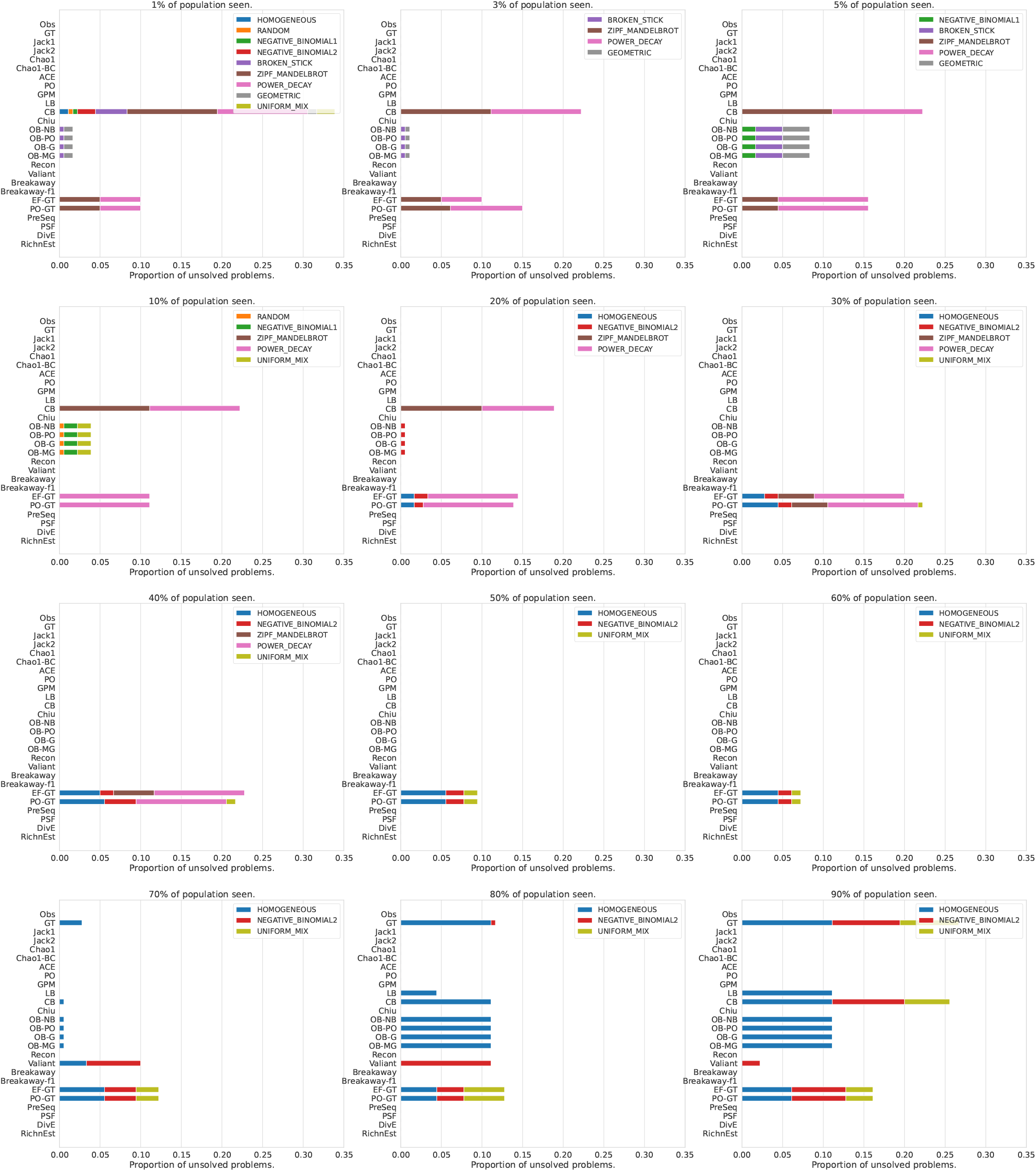
Proportion of unsolved problems for each method. Each subplot shows the proportion of unsolved problems for one subsampling rate. The colored bars further divide the unsolved problem by population model. The unsolvable problems vary for the tools and depend on both the subsampling rate and the population model

**Figure S.3:**
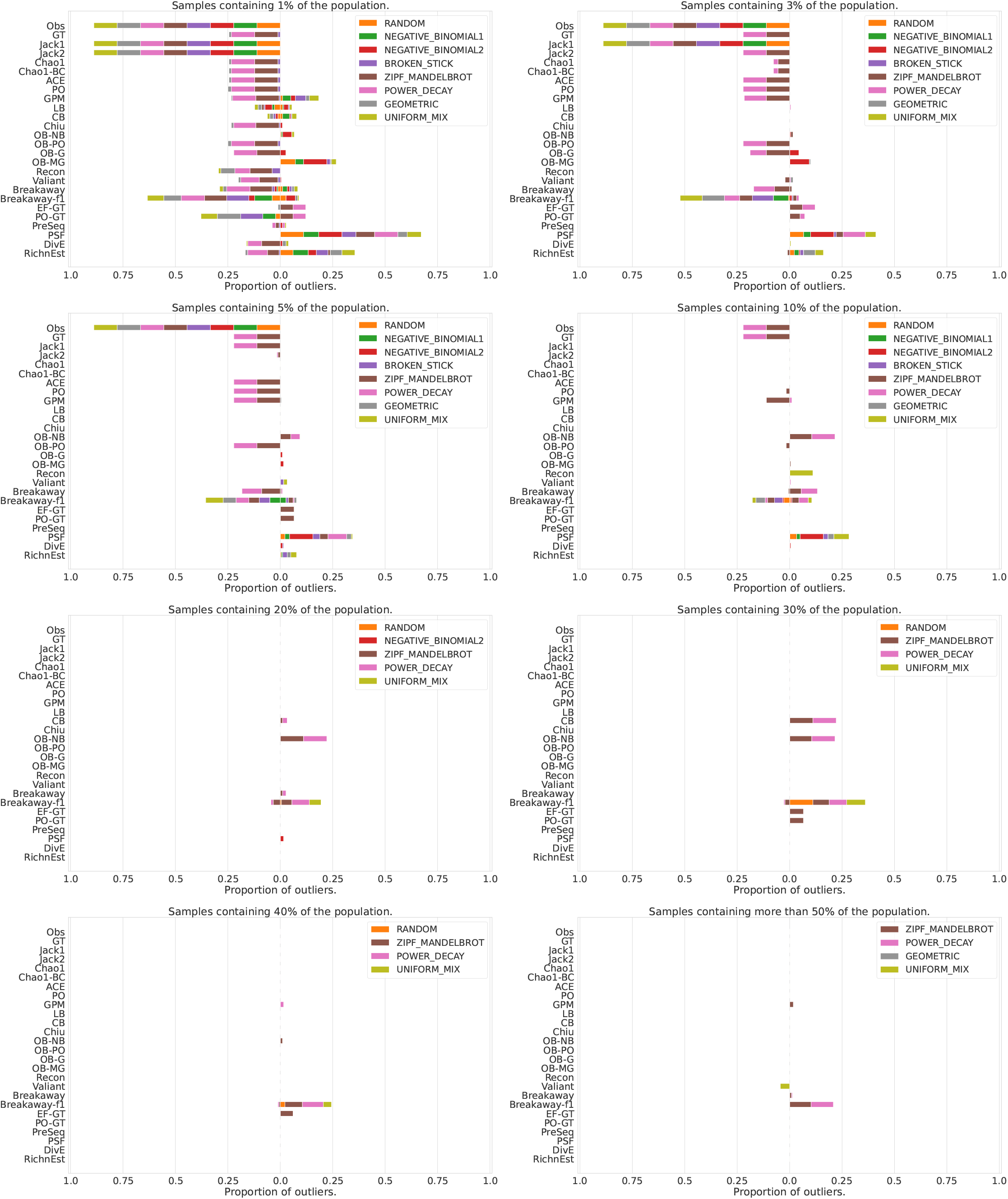
Proportion of outliers for each tool and each subsampling rate. The colored bars further divide the outliers by population model. To the left are the outliers that underestimate and to the right are the outliers that overestimate the true richness. An estimate is assumed to be an outlier if it is smaller than *S/c* or larger than *c* · *S*. For this plot, we used *c* = 2. The ‘hard’ problems with most outliers occur for subsampling rates below 0.1.

**Figure S.4:**
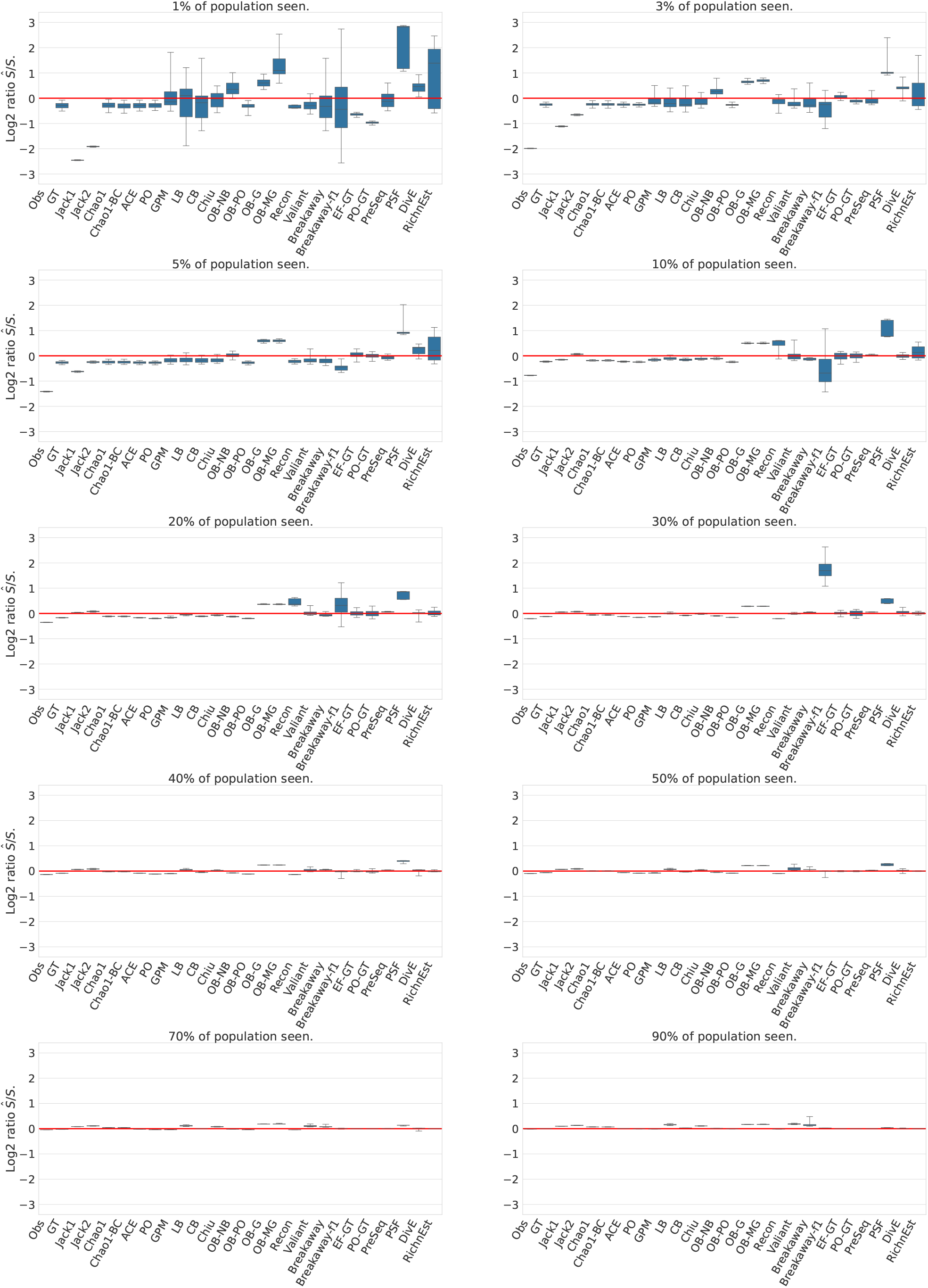
Estimation accuracy for the *random* population model. Boxplots display log_2_ ratios between the estimated species richness *Ŝ* and the true species richness *S*. If the prediction is correct, log_2_(*Ŝ/S*) = 0. If log_2_(*Ŝ/S*) *>* 0, the estimator overestimates the true richness and if log_2_(*Ŝ/S*) *<* 0, the estimator underestimates the true richness. Outliers have been removed (using a cutoff of log_2_(10)).

**Figure S.5:**
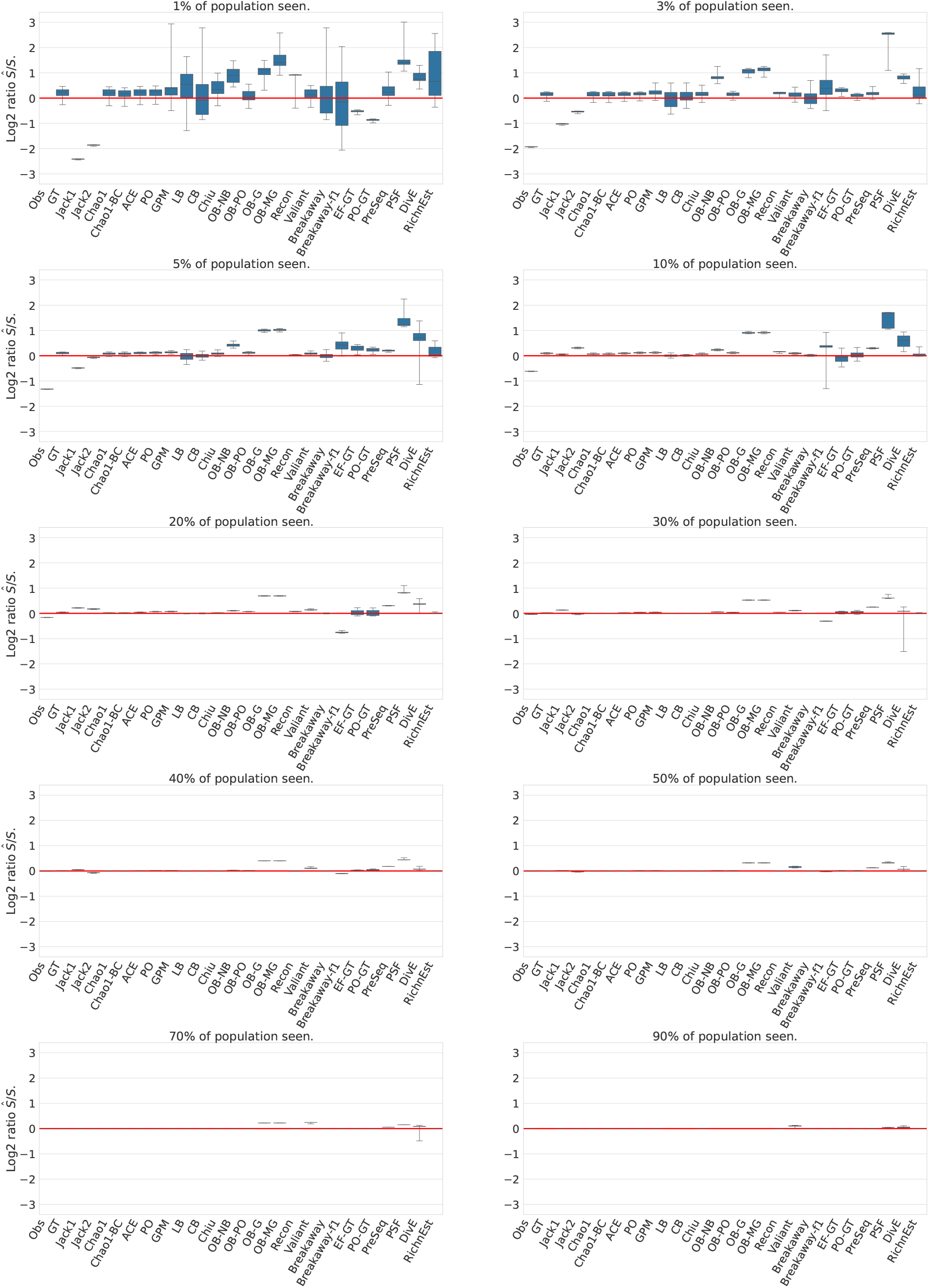
Estimation accuracy for the *homogeneous* population model. Boxplots display the log_2_ ratios between the predicted species richness *Ŝ* and the true species richness *S*. If the prediction is correct, log_2_(*Ŝ/S*) = 0. If log_2_(*Ŝ/S*) *>* 0, the estimator overestimates the true richness and if log_2_(*Ŝ/S*) *<* 0, the estimator underestimates the true richness. Outliers have been removed (using a cutoff of log_2_(10)).

**Figure S.6:**
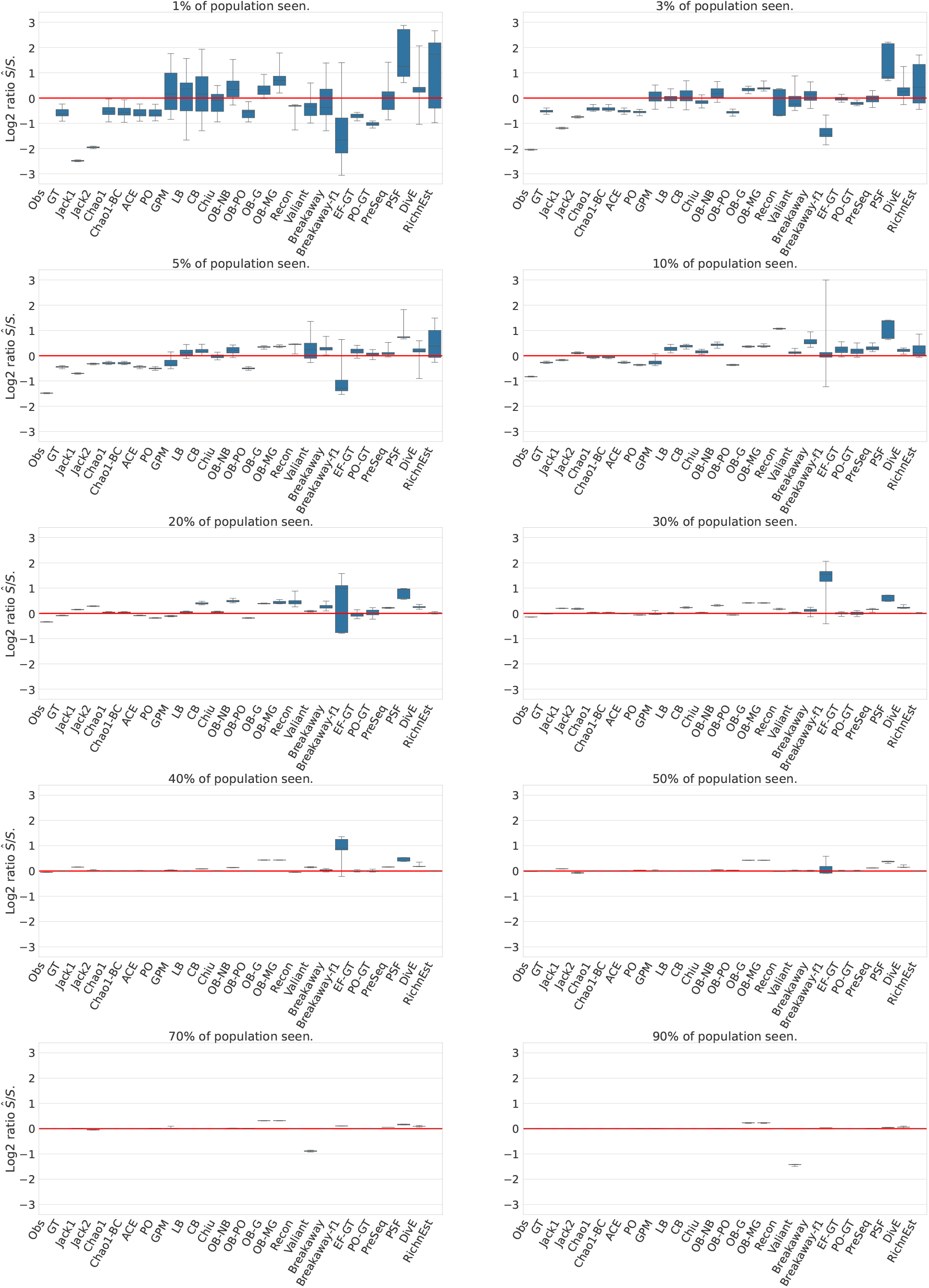
Estimation accuracy for the *uniform mix* population model. Boxplots display the log_2_ ratios between the predicted species richness *Ŝ* and the true species richness *S*. If the prediction is correct, log_2_(*Ŝ/S*) = 0. If log_2_(*Ŝ/S*) *>* 0, the estimator overestimates the true richness and if log_2_(*Ŝ/S*) *<* 0, the estimator underestimates the true richness. Outliers have been removed (using a cutoff of log_2_(10)).

**Figure S.7:**
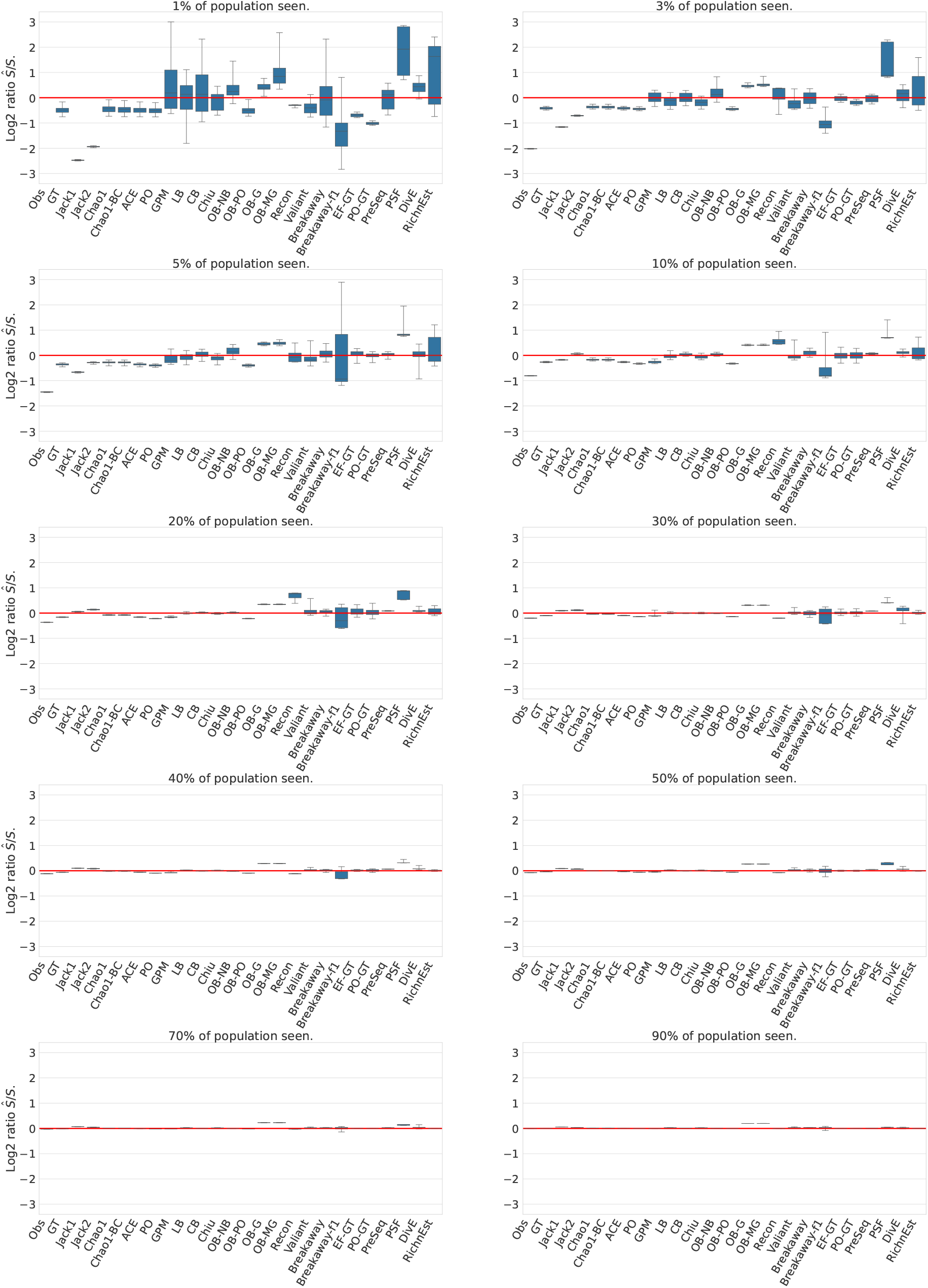
Estimation accuracy for the negative binomial population model (*r* = 2, *p* = 0.02). Boxplots display the log_2_ ratios between the predicted species richness *Ŝ* and the true species richness *S*. If the prediction is correct, log_2_(*Ŝ/S*) = 0. If log_2_(*Ŝ/S*) *>* 0, the estimator overestimates the true richness and if log_2_(*Ŝ/S*) *<* 0, the estimator underestimates the true richness. Outliers have been removed (using a cutoff of log_2_(10)).

**Figure S.8:**
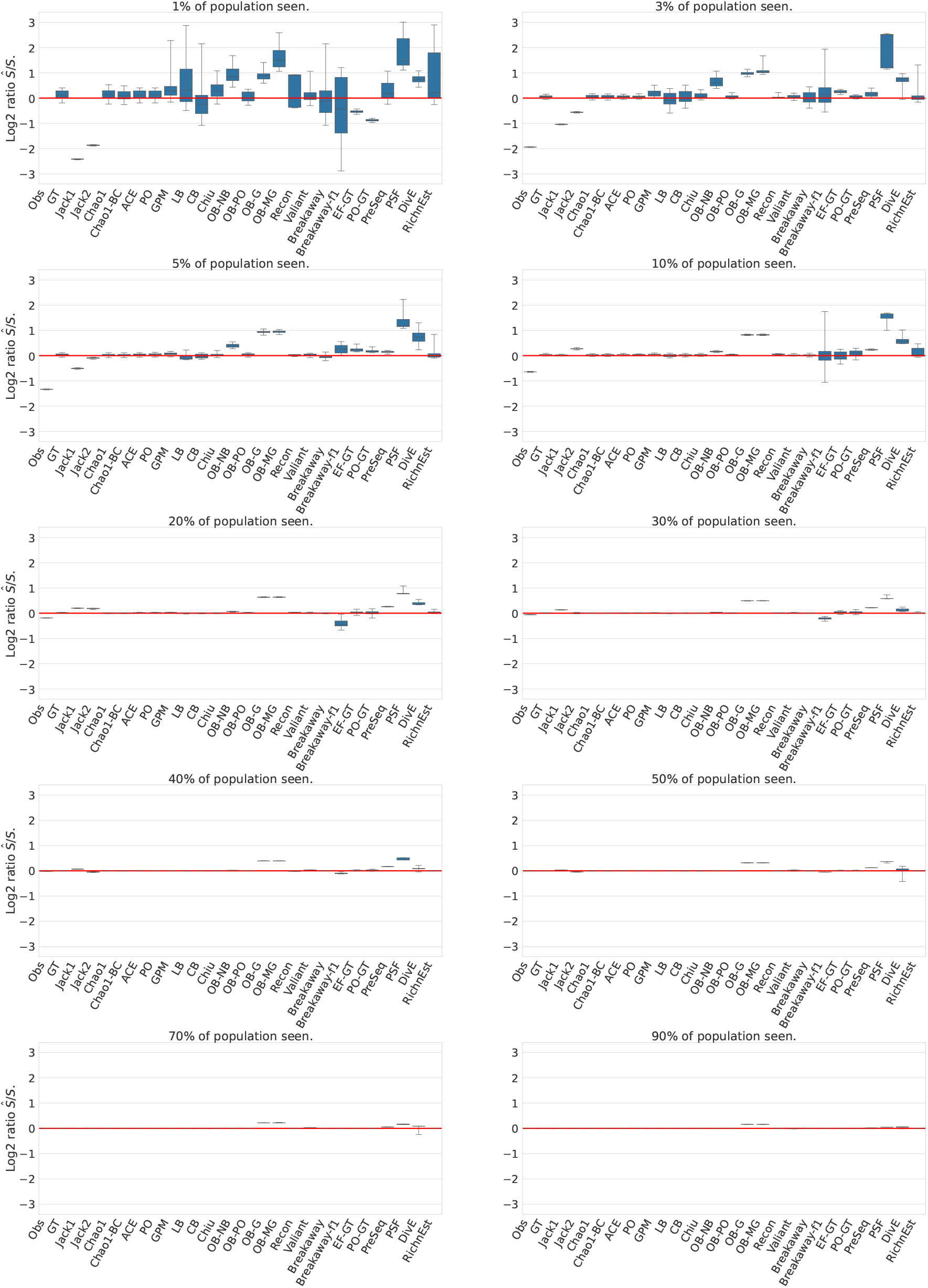
Estimation accuracy for the negative binomial population model (*r* = 20, *p* = 0.2). Boxplots display the log_2_ ratios between the predicted species richness *Ŝ* and the true species richness *S*. If the prediction is correct, log_2_(*Ŝ/S*) = 0. If log_2_(*Ŝ/S*) *>* 0, the estimator overestimates the true richness and if log_2_(*Ŝ/S*) *<* 0, the estimator underestimates the true richness. Outliers have been removed (using a cutoff of log_2_(10)).

**Figure S.9:**
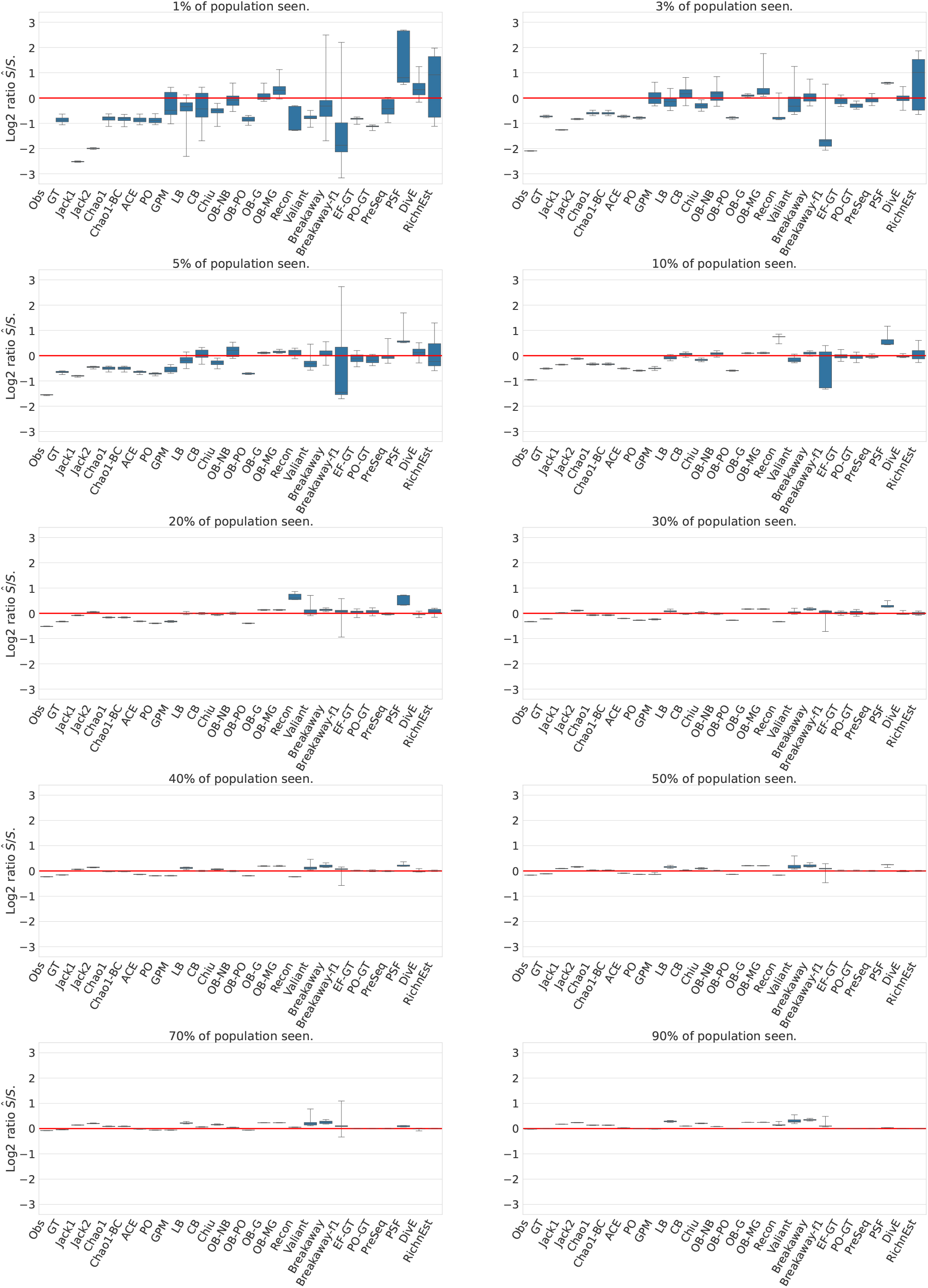
Estimation accuracy for the geometric population model (*p* = 1*/S*). Boxplots display the log_2_ ratios between the predicted species richness *Ŝ* and the true species richness *S*. If the prediction is correct, log_2_(*Ŝ/S*) = 0. If log_2_(*Ŝ/S*) *>* 0, the estimator overestimates the true richness and if log_2_(*Ŝ/S*) *<* 0, the estimator underestimates the true richness. Outliers have been removed (using a cutoff of log_2_(10)).

**Figure S.10:**
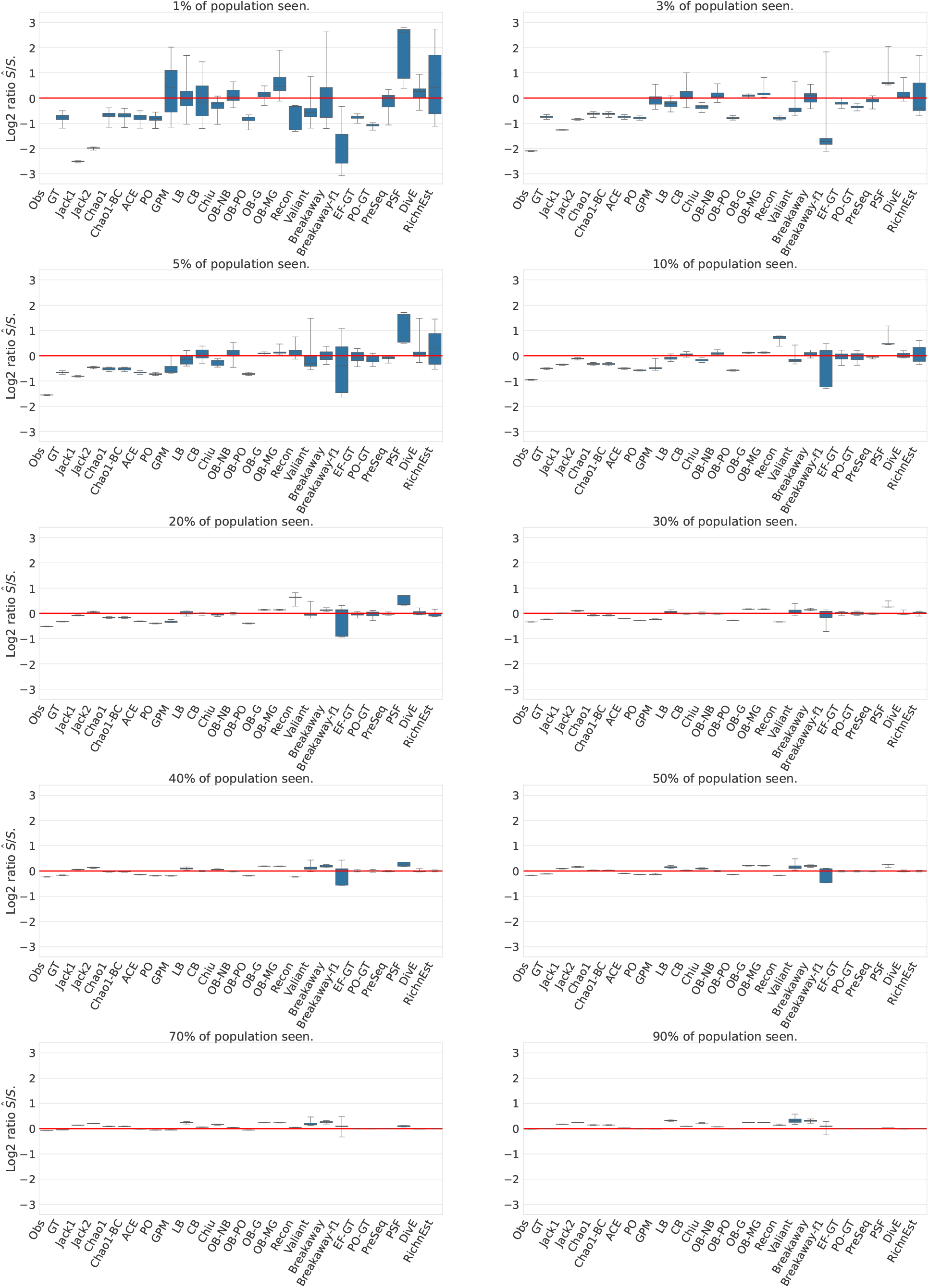
Estimation accuracy for the broken stick population model. Boxplots display the log_2_ ratios between the predicted species richness *Ŝ* and the true species richness *S*. If the prediction is correct, log_2_(*Ŝ/S*) = 0. If log_2_(*Ŝ/S*) *>* 0, the estimator overestimates the true richness and if log_2_(*Ŝ/S*) *<* 0, the estimator underestimates the true richness. Outliers have been removed (using a cutoff of log_2_(10)).

**Figure S.11:**
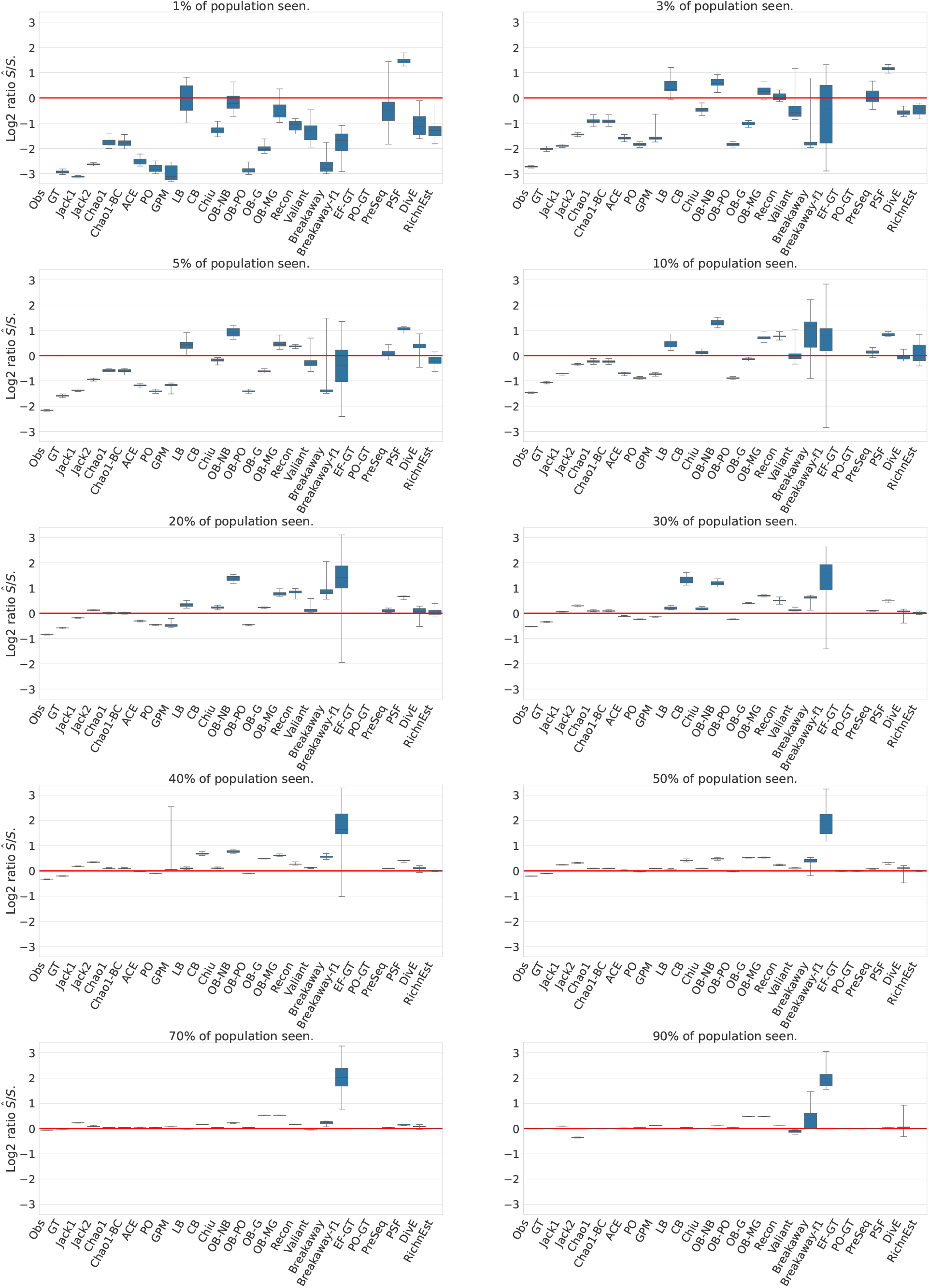
Estimation accuracy for the power decay population model. Boxplots display the log_2_ ratios between the predicted species richness *Ŝ* and the true species richness *S*. If the prediction is correct, log_2_(*Ŝ/S*) = 0. If log_2_(*Ŝ/S*) *>* 0, the estimator overestimates the true richness and if log_2_(*Ŝ/S*) *<* 0, the estimator underestimates the true richness. Outliers have been removed (using a cutoff of log_2_(10)).

**Figure S.12:**
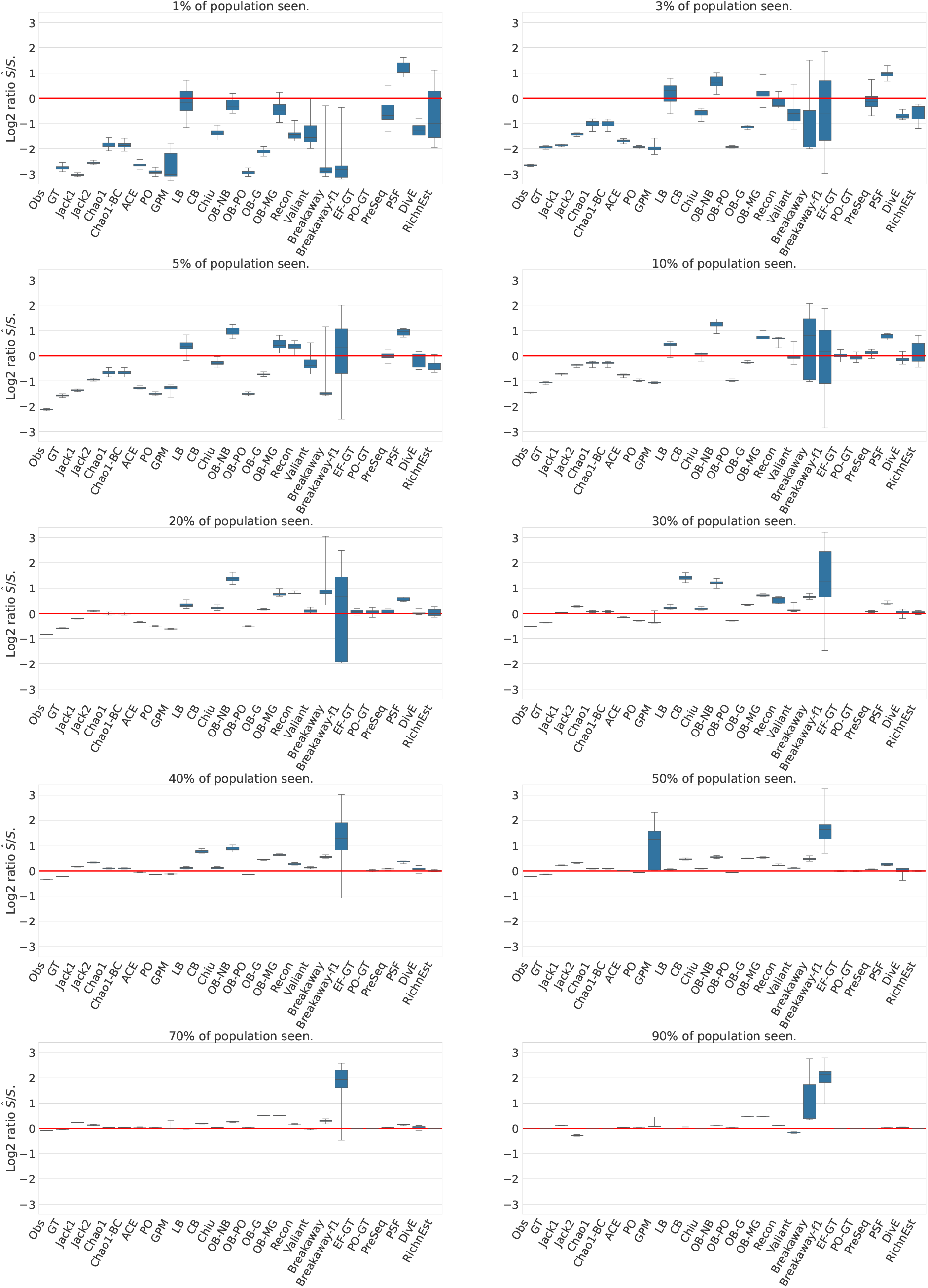
Estimation accuracy for the Zipf-Mandelbrot population model. Boxplots display the log_2_ ratios between the predicted species richness *Ŝ* and the true species richness *S*. If the prediction is correct, log_2_(*Ŝ/S*) = 0. If log_2_(*Ŝ/S*) *>* 0, the estimator overestimates the true richness and if log_2_(*Ŝ/S*) *<* 0, the estimator underestimates the true richness. Outliers have been removed (using a cutoff of log_2_(10)).

**Figure S.13:**
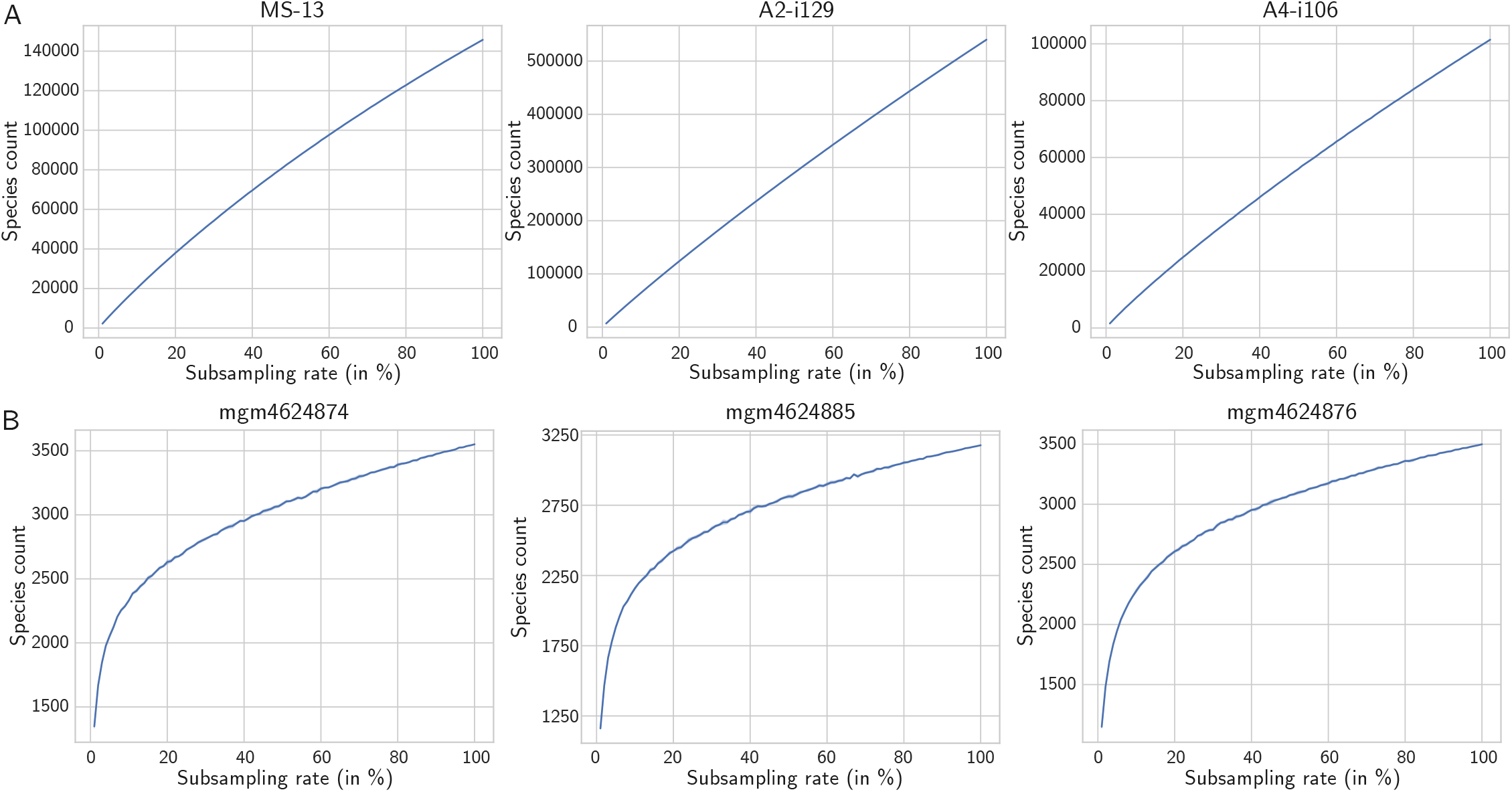
Rarefaction curves for sample richness (number of distinct species in the sample) for different subsampling rates (proportion of the sample included in the subsample), averaged over 100 subsamples. **A** 3 immune repertoire datasets (VDJTools Examples). **B** 3 microbiome datasets (MG-Rast database).

